# DIMPLE: A Tree-Guided Divide-and-Conquer Framework for ILS-Aware Phylogenetic Network Inference on Large-Scale Data

**DOI:** 10.64898/2026.10.12.758494

**Authors:** Ziyun Guang, Nicolae Sapoval, Zhi Yan, Mark Kessler, Luay Nakhleh

## Abstract

Inferring evolutionary histories of species provides a foundation for comparative genomics and evolutionary analyses. Extensive evidence of reticulate evolution motivates the inference of phylogenetic networks, which represent relationships that species trees cannot capture. However, accurate reconstruction of large phylogenetic networks remains a major challenge because of the vast space of possible network topologies. Here, we propose DIMPLE, a divide-and-conquer framework for phylogenetic network inference without an explicit restriction on network level. DIMPLE uses a tree-of-blobs (TOB), which represents the tree-like relationships between regions of potential reticulation while leaving relationships within those regions unresolved. Within each unresolved region, DIMPLE partitions the set of taxa into pairwise-disjoint subsets, each containing at most a user-specified number of taxa. Multiple division runs are allowed, and subsets from different runs may overlap. DIMPLE infers a subnetwork on each subset, merges the subnetworks within each region, and uses the TOB to assemble the resulting networks into a network on the full taxon set. Experiments with true inputs demonstrate accurate merging, with improved reticulation recovery over existing methods on higher-level networks. With estimated inputs, DIMPLE achieves competitive topological accuracy and more consistently recovers hybrid ancestry than competing methods across varying levels of network complexity. Its computational requirements support analyses of hundreds of taxa. Applications to fern and gymnosperm datasets recover documented hybridization and previously reported phylogenetic conflict.

## 1 Introduction

Phylogenetic trees have long served as the standard model of species evolution, but growing evidence shows that reticulate processes such as hybridization, introgression, and horizontal gene transfer cannot be adequately captured by a strictly bifurcating history (Doolittle 1999; Mallet et al. 2016). Rooted phylogenetic networks generalize trees to directed acyclic graphs in which internal nodes can have more than one parent, providing a more realistic representation of these events (Elworth et al. 2019). Accurate inference of such networks is now a central problem in phylogenomics. It remains computationally challenging: the number of possible tree topologies grows super-exponentially with the number of taxa, and allowing reticulations greatly expands this already vast search space.

Network inference must also account for incomplete lineage sorting (ILS), which can cause discordance among gene trees without reticulation. The multispecies network coalescent (MSNC) models both processes by describing how gene lineages trace their ancestry through a phylogenetic network. Under this model, calculating a gene tree’s probability requires accounting for the possible ancestral paths and coalescent histories that produce it (Yu et al. 2014), which makes likelihood-based calculations expensive for networks. Thus, existing likelihood-based inference methods remain limited to relatively small datasets (Wen et al. 2018; Elworth et al. 2019). Scalable likelihood-free alternatives also impose restrictions: for example, ALTS (Zhang et al. 2023) and Squirrel (Holtgrefe et al. 2025) do not model ILS, whereas NANUQ (Allman et al. 2019) targets level-1 network inference, and NetCS (Dai and Molloy 2026) and BROOQS (Arasti and Mirarab 2026) resolve an inferred tree-of-blobs (TOB) into a level-1 network. Scalable inference that accounts for both ILS and higher network complexity remains a challenge.

Divide-and-conquer is a general strategy for addressing large computational problems by solving smaller subproblems and combining their solutions. In phylogenomics, traditional divide-and-conquer approaches infer trees on overlapping subsets of taxa and combine them into a tree on the full taxon set using a supertree method. Shared taxa provide the information needed to relate the subset trees, as in SuperFine (Swenson et al. 2012) and FastRFS (Vachaspati and Warnow 2017). NJMerge (Molloy and Warnow 2019a) introduced a method for merging trees on pairwise-disjoint taxon subsets, using a distance matrix to relate taxa across subsets. It runs neighbor joining while enforcing topological constraints from the input trees. TreeMerge (Molloy and Warnow 2019b) subsequently improved robustness and scalability, whereas GTM (Smirnov and Warnow 2020) uses a guide tree and an unblended strategy that connects input trees without interleaving their internal structures.

Extending divide-and-conquer to phylogenetic networks presents additional challenges. First, reticulations may become non-identifiable when only a subset of taxa is considered. Division must therefore retain combinations of taxa that provide information about these events. Second, accurate subnetwork inference alone is insufficient to ensure accurate reconstruction of the full network: the merger must determine the relative positions of structures inferred on different subsets and identify representations of the same reticulation event across overlapping subnetworks. Inference errors can also cause subnetworks to imply conflicting evolutionary histories for the same taxa. The merger must reconcile these conflicts and assess whether the retained information is sufficient to reconstruct the full network.

Existing approaches address parts of these challenges. NetMerger (Zhu et al. 2019) combines subnetworks inferred on triples of taxa, but the cubic number of triples creates a computational bottleneck. NANUQ^+^ (Allman et al. 2025) takes a TOB-guided divide-and-conquer approach, resolving individual multifurcations of the inferred TOB to infer a level-1 network. InPhyNet (Kolbow et al. 2026) uses distance information to merge larger subnetworks, although its input subnetworks are restricted to level 1, which can in turn limit its ability to recover higher-level reticulation structure.

These challenges motivate using a guide that helps division retain reticulation information and helps merging place the inferred structures relative to one another. Prior approaches to species tree estimation, including TreeMerge (Molloy and Warnow 2019b) and GTM (Smirnov and Warnow 2020), use trees on the full taxon set, such as those inferred with ASTRAL (Zhang et al. 2018), to guide division or merging. Extending this strategy to phylogenomic network inference requires considering which features of the network a guide tree can consistently recover. Dinh and Baños (2025) showed that an optimal solution to the weighted quartet consensus (WQC) problem, the criterion used by ASTRAL (Zhang et al. 2018), need not converge to a displayed tree of the underlying network. A TOB provides an alternative guide and is already an established starting point for scalable network inference. It summarizes the tree-like relationships between regions containing reticulation events, called blobs, while leaving their internal structure unresolved. The TOB can be estimated from gene-tree quartet information using TINNiK (Allman et al. 2024) or TOB-QMC, implemented in TREE-QMC (Dai et al. 2026). DIMPLE builds on this established use of the TOB to guide division and network assembly, using subnetwork inference and merging to reconstruct blobs that may contain multiple reticulations.

We introduce **DIMPLE**: a **D**ivide-**I**nfer-**M**erge framework for **P**hylogenetic network inference on **L**arge-scal**E** data, with no explicit upper bound on the level of the final network. DIMPLE divides taxa into subsets using an estimated TOB, infers a subnetwork on each subset using the maximum pseudo-likelihood (MPL) criterion (Yu and Nakhleh 2015) implemented in PhyloNet (Wen et al. 2018), and merges the subnetworks into a network on the full taxon set. The subnetwork-inference step uses an existing method; our contributions are the division and merger algorithms. The decomposition step forms taxon subsets according to the TOB structure and repeats this process across runs to increase reticulation coverage. Inspired by InPhyNet, our merger combines the resulting potentially overlapping subnetworks by constructing a distance-based neighbor-joining backbone guided by representative displayed trees selected from the subnetworks. It then evaluates candidate reticulation placements using pseudo-likelihood, retaining insertions that meet the acceptance threshold. Finally, the reconstructed regions are assembled according to the TOB using an unblended strategy similar to GTM.

We evaluate DIMPLE on simulated networks with 50, 100, and 250 taxa under the MSNC model. DIMPLE has zero topological distance in a substantial fraction of cases and improves upon competing methods in several higher-level network settings. We also apply DIMPLE to the fern and gymnosperm datasets from the One Thousand Plant Transcriptomes (1KP) project. The inferred networks recover the documented allopolyploid origin of *Polypodium hesperium* and represent previously reported conflict concerning the placement of Gnetales relative to conifers.

## 2 Materials and Methods

### 2.1 Background

A *phylogenetic network N* on a taxon set *X* is a rooted directed acyclic graph with vertex set *V* (*N*) and edge set *E*(*N*). The vertex set consists of four types of nodes: a single *root node ρ* with in-degree 0; a set of *leaves V*_*L*_ with out-degree 0, bijectively labeled by the taxa in *X*; a set of internal *reticulation nodes V*_*R*_ with in-degree greater than 1; and a set of internal *tree nodes V*_*T*_ with in-degree 1. This work focuses on *binary* phylogenetic networks, which satisfy the following additional constraints: the root *ρ* has out-degree 2; every tree node *v* ∈ *V*_*T*_ has out-degree 2; every reticulation node *v* ∈ *V*_*R*_ has in-degree 2 and out-degree 1; and every leaf *v* ∈ *V*_*L*_ has in-degree 1. A *reticulation edge* is an edge whose head is a reticulation node; all other edges are *tree edges*.

The *level* of a phylogenetic network is the maximum number of reticulation nodes in any biconnected component of its underlying undirected graph. The *restriction* of *N* to *Y* ⊆ *X*, denoted *N*| _*Y*_, is formed by computing the homeomorphic subnetwork of *N* induced on *Y* and suppressing nodes of in-degree and out-degree one. A *phylogenetic tree* is a phylogenetic network without reticulation nodes. A *displayed tree* of *N* is a tree that can be obtained by removing at each reticulation node all but one of the reticulation edges with the same target node and suppressing the resulting degree-2 vertices.

The TOB summarizes the tree-like relationships between regions of a phylogenetic network containing reticulation events. Each such region, called a *blob*, is contracted into a single vertex, hiding its internal reticulation structure while retaining its connections to the rest of the network (Dai et al. 2026; Dai and Molloy 2026). For example, contracting each colored blob in Fig. 1 produces the TOB. Formally, blobs are maximal bridgeless subgraphs of the underlying undirected graph. A blob is *nontrivial* if it contains more than one vertex. The TOB, denoted *T*_*N*_, is obtained by contracting the edges within each blob and suppressing vertices with in-degree 1 and out-degree 1 and any root with only one child.

**Figure 1.**
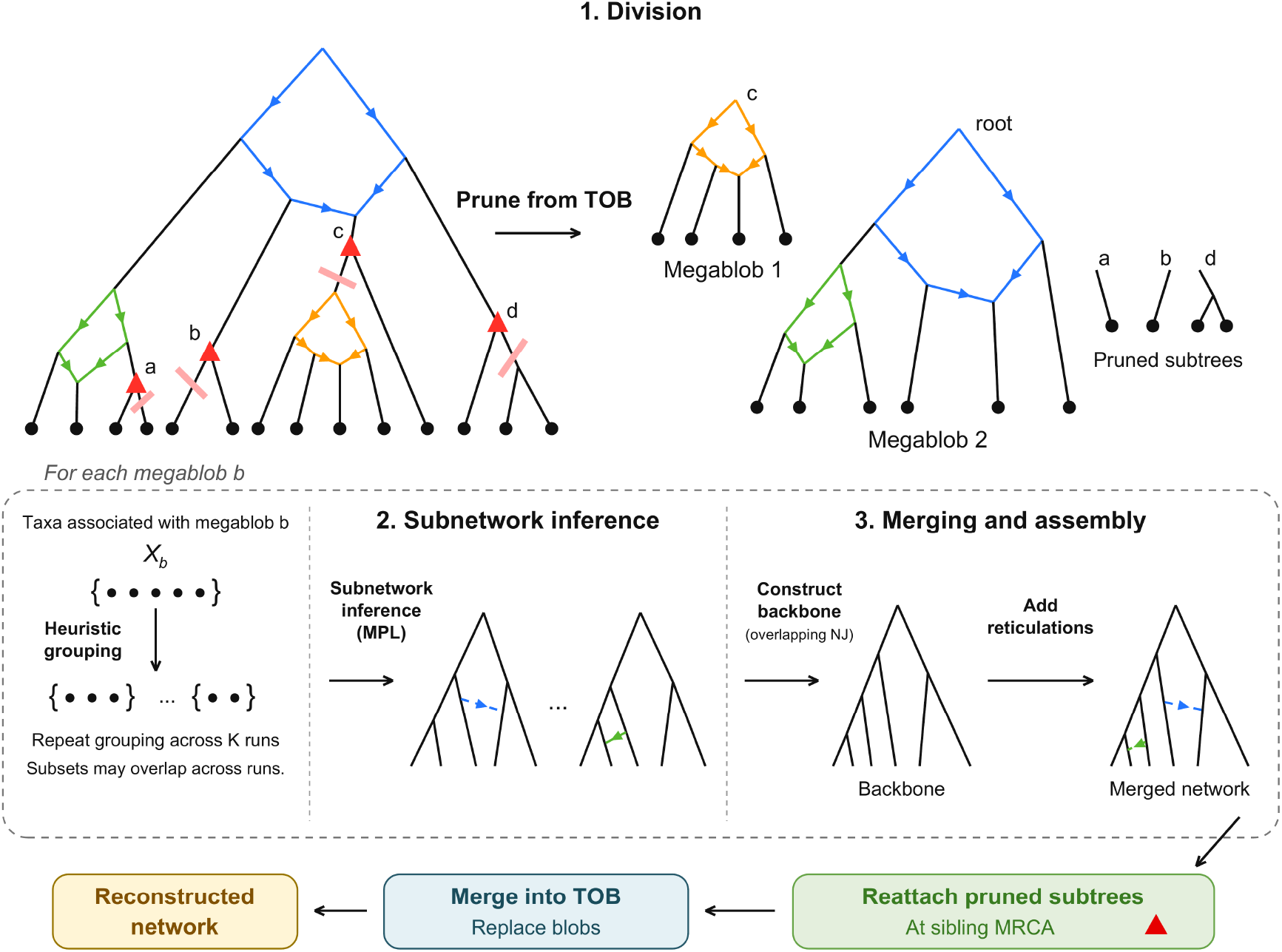
Pipeline of the DIMPLE method. Colored blobs are shown expanded to illustrate their boundaries; their internal structures are unknown during inference. First, megablobs are identified from the TOB, and subtrees outside them are pruned off. For each megablob, heuristic grouping is applied to partition the set of leaves into subsets. A subnetwork is inferred for each subset using MPL; these subnetworks serve as inputs to the merger. After the merger returns a merged network per megablob, the pruned subtrees are reattached and each megablob is expanded back into the TOB to form the final network.

A tree *refines T*_*N*_ if *T*_*N*_ can be obtained by contracting edges in that tree. We also define a *megablob* as a connected component of the subgraph of *T*_*N*_ induced by vertices representing nontrivial blobs. Thus, blobs represented by adjacent vertices in *T*_*N*_ belong to the same megablob and are processed jointly by our method.

### 2.2 DIMPLE framework

Given a collection of gene trees *G* = {*g*_1_, …, *g*_*q*_} on a taxon set *X*, our goal is to infer a rooted phylogenetic network 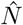 on *X* under the MSNC model. DIMPLE uses a TOB inferred from these gene trees to guide reconstruction.

DIMPLE reconstructs the network in three steps (Fig. 1). First, guided by the TOB, it partitions the set of taxa associated with each megablob into pairwise-disjoint subsets, each containing at most a user-specified number *s* of taxa. For each megablob with more than *s* taxa remaining, it performs *K* grouping attempts. The subsets within each partition are pairwise disjoint, whereas subsets from different partitions may overlap. Second, it infers a subnetwork on each subset using the MPL criterion (Yu and Nakhleh 2015) implemented in PhyloNet (Wen et al. 2018). Third, it merges the inferred subnetworks within each megablob, reattaches the pruned subtrees, and assembles the resulting networks according to the TOB to obtain 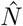 on the full taxon set *X*.

### 2.3 Merger problem formulation

The merger combines subnetworks inferred for each megablob before assembling the full network according to the TOB. Throughout this section, *b* denotes a megablob of the inferred TOB 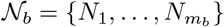 be the collection of *m*_*b*_ subnetworks inferred across the division runs for *b*. Write *X*_*i*_ = *L*(*N*_*i*_) for the taxon set of subnetwork *N*_*i*_ and 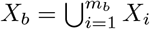 for their union. Let *G*_*b*_ denote the input gene trees restricted to *X*_*b*_. We formulate the task of merging these subnetworks into a rooted network on *X*_*b*_.

#### Definition 1

(Pairwise compatibility). A collection of subnetworks N_*b*_ is *pairwise compatible* if, for every

*i* ≠ *j* with *X*_*i*_ ∩ *X*_*j*_ ≠ ∅,

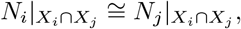

where ≅ denotes an isomorphism preserving the root and taxon labels.

#### Definition 2

(Compatibility supernetwork). A rooted phylogenetic network 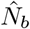 on *X*_*b*_ is a *compatibility supernetwork* for N_*b*_ if 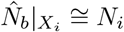 for every *i* = 1, …, *m*_*b*_.

Pairwise compatibility describes agreement on overlaps but does not guarantee the existence of a compatibility supernetwork.

#### Definition 3

(Ideal blob merger). Given N_*b*_ that admits a compatibility supernetwork and gene trees G_*b*_, find a compatibility supernetwork 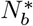 that maximizes the pseudo-likelihood given G_*b*_.

In practice, subnetworks are inferred independently and may disagree on shared taxa, preventing a compatibility supernetwork from existing. We therefore also consider the following objective.

#### Definition 4

(Maximum-compatible-subset merger). Given *N*_*b*_ and *G*_*b*_, find a subset *C*^⋆^ ⊆ *N*_*b*_ and a rooted phylogenetic network 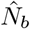 bon *X*_*b*_ such that:

1. 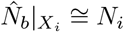 for every *N*_*i*_ ∈ C;
2. each reticulation of 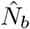 represents an event present in at least one subnetwork in *C*^⋆^, with no event represented more than once in 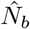;
3. *C*^⋆^ has maximum cardinality among subsets of *N*_*b*_ for which there exists a rooted phylogenetic network on *X*_*b*_ satisfying conditions (1) and (2); and
4. for the selected 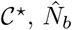 maximizes the pseudo-likelihood given _*b*_ among networks satisfying conditions (1) and (2).

The maximum-compatible-subset merger defines an ideal objective for potentially conflicting inputs. Our implementation uses a heuristic: DIMPLE selects a maximum-cardinality pairwise-compatible collection of representative displayed trees to guide backbone construction, relaxing constraints when necessary.

### 2.4 DIMPLE algorithms

#### 2.4.1 Decomposition

Decomposition aims to retain information about reticulation events while limiting each subnetwork inference problem to at most *s* taxa. Let *v* be a vertex of the inferred TOB representing a nontrivial blob, let *d* be the number of outgoing edges of *v*, denote these edges by *c*_1_, …, *c*_*d*_, and let *X*(*c*_*j*_) be the taxon set below *c*_*j*_. These taxon sets are represented by solid circles in the diagrams labeled “Megablob 1” and “Megablob 2” in Fig. 1. The TOB records these descendant clades but leaves their relationships within that blob unresolved.

We group taxa from several clades *X*(*c*_*j*_) into the same subset to provide information about these relationships. Ideally, a subset would contain at least one taxon from every *X*(*c*_*j*_), but this is impossible when *d > s*. We therefore prune descendant subtrees first and group the remaining taxa into subsets of size at most *s*. This strategy aims to retain information about reticulations within subsets but does not guarantee recovery of every reticulation.

Directly adjacent blob vertices are processed together as megablobs. For each megablob, we isolate the corresponding component of the TOB and apply the following two steps (Fig. 1):

1. Pruning. While the component contains more than *s* taxa, we greedily remove eligible descendant subtrees. We do not cut edges leaving blob vertices, and each cut must leave at least three taxa on each side. Pruning stops when at most *s* taxa remain or no eligible cut exists. The pruned subtrees are retained for reattachment.
2. Heuristic grouping. We treat the remaining subtrees below edges leaving the megablob as items, each weighted by its number of taxa, and split items larger than *s* before grouping. If their total weight is at most *s*, they form one subset. Otherwise, the first attempt greedily forms the first group from the smallest items and randomly packs the remaining items. Subsequent attempts randomly pack all items subject to capacity *s*. We perform *K* attempts and discard duplicate partitions.

Within each partition, the taxon subsets are pairwise disjoint; subsets from different partitions may overlap. Repeated grouping provides additional opportunities to retain information about a reticulation within a single subset. TOB inference using TOB-QMC (Dai et al. 2026) takes *O*(*qn*^3^) time, where *n* = |*X*| and *q* is the number of input gene trees. For a fixed number of division attempts, pruning and grouping require *O*(*n*^2^) time in the worst case.

#### 2.4.2 Merging

For each megablob *b*, DIMPLE constructs a tree backbone using representative displayed trees as constraints, then inserts candidate reticulations extracted from the subnetworks. After the pruned subtrees are reattached, the resulting networks are assembled according to the structure of the TOB.

##### Constraint tree selection

For each subnetwork *N*_*i*_∈ N _*b*_, we enumerate its displayed trees and select one using rooted-triplet support from the gene trees. Let *T* (*N*_*i*_) denote the set of these displayed trees and rt(*T*) the set of rooted triplets displayed by a tree *T*. For a rooted triplet *t*, define its gene-tree support as

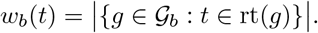

Let

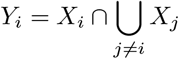

be the taxa shared with other subnetworks of *b*. We select

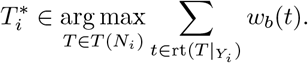

If fewer than three taxa are shared, all candidate trees receive a score of zero. The selected trees serve as candidate constraints for backbone construction. Triplet counts are computed once and reused, while displayed-tree enumeration is exponential in the number of reticulations per input subnetwork.

##### Compatible-tree selection

The selected displayed trees may disagree on shared taxa. We construct a compatibility graph with one vertex for each selected tree and an edge between two vertices if their trees agree as unrooted trees on their shared taxa. Trees with disjoint taxon sets are also connected. A maximum clique identifies a maximum-cardinality pairwise-compatible collection of trees, which we retain as constraints for backbone construction. Maximum-clique selection has exponential worst-case complexity in the number of candidate constraint trees.

##### Overlapping neighbor joining

We construct the backbone 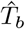 using a neighbor-joining procedure inspired by NJMerge (Molloy and Warnow 2019a). For each pair of taxa, the initial distance is the mean path length across input gene trees containing both taxa, where path length is the sum of branch lengths along the path connecting the two taxa. Let *A* be the set of current tree labels, *m* = |*A*|, and *D* the current distance matrix.

For distinct labels *a, a*^′^ ∈ *A*, the neighbor-joining criterion is

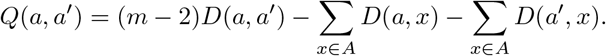

At each iteration, we consider pairs in increasing order of *Q* and select the first pair whose labels are siblings in every retained constraint tree containing both labels. If no pair satisfies this condition, we relax the constraints and select the minimum-*Q* pair. We join the selected pair and update the distances and constraint trees, repeating until a single backbone remains. With a fixed number of constraint trees, backbone construction takes 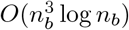 time, where *n*_*b*_ = |*X*_*b*_|.

##### Greedy reticulation insertion

We extract candidate reticulations from the inferred subnetworks and group candidates identified as representing the same event. We process insertion candidates in dependency order: a candidate is given priority over another when its descendant taxa overlap the taxa below the other child of the parent whose connection will be added for the second candidate. For each candidate, we enumerate insertion locations consistent with the source subnetwork information. We evaluate each possible reticulation placement using the log pseudo-likelihood under the MSNC, with all branch lengths and inheritance probabilities held fixed. A candidate is accepted at its highest-scoring placement only if it improves the current network’s score. More details are described in the appendix.

##### Reattachment and assembly

After merging the subnetworks for each megablob *b*, we reattach the subtrees removed during decomposition to 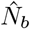, using *T*_*N*_ to guide their placement. We then replace each megablob in *T*_*N*_ with its expanded network, retaining the surrounding tree-like structure, to obtain the full network 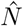 on *X*. The computational cost grows with the number of candidate reticulations, the number of insertion locations, and the cost of each pseudo-likelihood evaluation. A detailed runtime analysis is provided in Appendix C.

## 3 Evaluation

### 3.1 Simulated data

We simulated networks with 50, 100, and 250 taxa under the birth-death-hybridization (BDH) model using SiPhyNetwork v1.1.0 (Justison et al. 2023) in R 4.5.3. We selected 10 networks each at levels 1, 2, and 3+ for each dataset size, giving 90 networks in total. To simulate the networks, we set the rates of lineage-generative, lineage-degenerative, and lineage-neutral reticulations to 0.5, 0.25, and 0.25, respectively. The inheritance probability was drawn from a Beta(10,10) distribution. We used a birth rate of 1, a death rate of 0.1, and hybridization rates of 0.025, 0.05, and 0.1. For 50-taxon networks, we additionally varied the simulation parameters and increased the maximum hybridization rate to 0.27 to obtain enough higher-level networks. Finally, we added an outgroup taxon as sister to the simulated ingroup. The reported numbers of taxa exclude the outgroup.

Each network was evaluated under low- and high-ILS conditions. For 50-taxon networks, we multiplied network branch lengths by 2 and 0.5 to generate the low- and high-ILS conditions, respectively. For 100- and 250-taxon networks, the unscaled simulated networks already exhibited high ILS, so we multiplied branch lengths by 4 to generate the low-ILS condition. We quantified ILS levels using species trees simulated under the corresponding settings, computing the unrooted normalized Robinson–Foulds (RF) distance between each species tree and the gene trees simulated within it. These distances were approximately 26% under low ILS and 62% under high ILS across all three dataset sizes.

For each network, we simulated 1,000 gene trees under the MSNC using the simGTinNetwork command in PhyloNet v3.8.2 (Wen et al. 2018). We converted gene-tree branch lengths from coalescent units to expected substitutions per site using a scaling factor of 0.018. Along each gene tree, we simulated a 500-bp sequence alignment using INDELible v1.03 (Fletcher and Yang 2009) under a Hasegawa–Kishino–Yano (HKY) substitution model (*κ* = 1; base frequencies *π*_*T*_ = 0.2112, *π*_*C*_ = 0.2888, *π*_*A*_ = 0.2896, and *π*_*G*_ = 0.2104), with continuous gamma-distributed rate heterogeneity (shape parameter 1.8) and no invariant sites. Insertions and deletions were generated under a power-law length distribution (a = 1.8, maximum length 16) at a rate of 0.1 relative to the substitution rate. We realigned the sequences for each gene using MAFFT v7.526 (Katoh and Standley 2013) and inferred gene trees from the resulting alignments using IQ-TREE v3.0.1 (Wong et al. 2026).

### 3.2 Biological data

To evaluate DIMPLE on biological data, we analyzed the fern and gymnosperm subsets of the 1KP dataset (One Thousand Plant Transcriptomes Initiative 2019), previously used to evaluate InPhyNet (Kolbow et al. 2026) and TOB-QMC (Dai et al. 2026). The fern dataset includes lineages with documented hybridization and polyploidy, including *Polypodium* (Sigel et al. 2014). The gymnosperm dataset includes Gnetales, whose relationship to conifers has received conflicting support across phylogenetic studies (Bowe et al. 2000; Ran et al. 2018; Stull et al. 2021).

We applied DIMPLE, InPhyNet, CAMUS (Willson and Warnow 2026), NetCS, and BROOQS to these biological datasets using the settings described in Section 3.3. Unlike the simulated data, these datasets have no known ground-truth network. We therefore compare the inferred relationships with published biological evidence.

### 3.3 Parameter settings and methods evaluated

#### 3.3.1 TOB estimation

For each simulated network, we inferred the TOB from the estimated gene trees using TOB-QMC with significance thresholds *α* = 10^−7^ and *β* = 0.95. For the biological analyses, we used the same TOB-QMC thresholds. We rooted the estimated gene trees, ASTRAL trees, and inferred TOBs using the designated outgroup.

#### 3.3.2 DIMPLE division and subnetwork inference

During the division step, we set our maximum subset size to 12, which is a reasonable number that allows efficient computation by network inference methods. We selected *K* = 5 division runs based on the tradeoff between accuracy and computational cost examined in Appendix G (Figs. S2, S3, and S4); all DIMPLE results in the main text use this setting. We also tested maximum subset sizes of 6 and 18 on the 100-taxon dataset (Appendix H; Fig. S5).

For each subset of leaves, we inferred a subnetwork from gene trees restricted to that subset. We ran MPL using PhyloNet’s `InferNetwork_MPL` command by setting MPL’s maximum allowed number of reticulations to the number of reticulations *h* in the corresponding ground-truth subnetwork. Supplying the true *h* removes the model-selection error that would arise from estimating *h*. More realistically, estimating *h* would require running with *h* = 0, 1, 2, … and picking the best via a selection criterion. As estimating *h* is not our main focus here, the reported accuracy isolates the performance of the divide-and-merge pipeline itself without the confounding of *h*-selection error. During MPL inference, we chose 10 search runs and selected the network with the best pseudo-likelihood score. We used the ASTRAL-inferred tree restricted to each subset of taxa as a starting tree to guide MPL search for each subnetwork.

For the biological data, the number of reticulations was selected manually. Within each blob subnetwork, we ran MPL with maximum allowed numbers of reticulations *h* = 0, 1, 2, 3 and recorded the highest log pseudo-likelihood attained at each *h*. Following the discussion of score improvements as reticulations are added in Cao et al. (2023, Section 5.4), we used the resulting score profiles to assess qualitatively the number of reticulations supported by each subnetwork (Appendix I; Fig. S6).

#### 3.3.3 Comparison methods and parameter settings

We compared the DIMPLE merger with the InPhyNet merger and evaluated the complete DIMPLE pipeline against InPhyNet, CAMUS, NetCS, and BROOQS on simulated datasets with 50, 100, and 250 taxa. Although the last three methods do not use divide-and-conquer approaches and are not designed for higher-level networks, they provide additional comparisons as methods designed for scalable network inference.

For InPhyNet’s subset decomposition, we followed the InPhyNet implementation in three parts: first, we ordered the taxa by a neighbor-joining tree of the average gene-tree internode distance (AGID) matrix and partitioned them into overlapping contiguous blocks of at most 50 taxa; next, we ran TINNiK (*α* = 0.01, *β* = 0.99) on each block to estimate its TOB; finally, we selected one representative leaf per blob, collapsed each blob and preserved only a single representative on the ASTRAL species tree, applied centroid-edge (SATe) decomposition to the collapsed tree, and re-expanded the blobs so that all taxa incident to a common reticulation cycle remain within a single subset. For InPhyNet’s subnetwork inference using SNaQ (Solís-Lemus and Ané 2016), we also gave SNaQ the true number of reticulations obtained by restricting the ground-truth network to each leaf set. For CAMUS, we also selected the network whose number of reticulations equals the number in the ground-truth network to represent a best-case selection; for the few replicates in which the true number exceeded the largest that CAMUS produced, we selected the network with the closest available number of reticulations.

For the biological datasets, we ran InPhyNet using the taxon subsets reported in its supplementary information, and SNaQ was run at the reticulation count reported for each subset. For CAMUS, we examined the improvement in its optimization score as reticulations were added, following the authors’ recommendation. Further details are given in Appendix I.

### 3.4 Simulation experiments

#### Experiment 1: Reticulation retention during division

We evaluate division on simulated data by counting the ground-truth reticulations retained and measuring level loss within the resulting subsets. A reticulation is preserved when both of its parent lineages and the hybrid are retained within a single subset. We report level loss as the level of the ground-truth network minus the maximum level among subnetworks on the taxon subsets produced by the divider. This measures whether reticulations remain together within one subnetwork. For example, division may separate the two reticulations of a level-2 blob into different subsets, each inducing a level-1 network. Both reticulations are retained across the subsets, giving zero reticulation loss, but the level loss is one.

Specifically, we compare four division strategies. **TOB no restriction** separates the tree-like regions of the inferred TOB while leaving each blob intact, without imposing a subset-size limit. It provides a reference before further division into size-constrained subsets. **DIMPLE** uses the inferred TOB with our default 5 division attempts. **InPhyNet TINNiK** uses InPhyNet’s decomposition algorithm, whereas **InPhyNet NJ** uses InPhyNet’s naive NJ-tree clade-cutting approach.

#### Experiment 2: Network reconstruction with true inputs

To evaluate merger performance without TOB or subnetwork estimation error, we apply DIMPLE’s division to the true TOB collapsed from the ground-truth network and obtain true subnetworks by restricting the ground-truth network to the resulting taxon subsets. DIMPLE and the InPhyNet merger both receive true subnetworks obtained using DIMPLE’s divider. The InPhyNet merger uses subnetworks from a single partition because it requires pairwise-disjoint taxon sets, whereas DIMPLE combines subnetworks across division runs, whose taxon sets may overlap. Although NetCS and BROOQS do not merge subnetworks, we include them as comparisons using the true TOB and true simulated gene trees for a controlled study of the mergers. NetCS (Dai and Molloy 2026) and BROOQS (Arasti and Mirarab 2026) have shown high accuracy when given the true TOB. NetCS does not resolve 4-blobs, but all cycles in our simulated networks contain at least five vertices. We started with 10 networks for each combination of dataset size, network level, and ILS condition.

#### Experiment 3: End-to-end inference with estimated inputs

We compare the complete DIMPLE pipeline with InPhyNet, CAMUS, NetCS, and BROOQS on all simulated networks, starting from the same 1,000 estimated gene trees per network. Each method uses guide trees inferred from these gene trees. ASTRAL is also included for comparison.

### 3.5 Evaluation criteria

#### Gene tree inference error and guide tree accuracy

We assess the accuracy of the estimated gene trees and the two guide trees used by the inference methods: the TOB and the ASTRAL species tree. Gene tree estimation error is the unrooted normalized RF distance between estimated and true gene trees, averaged over the 1,000 genes per simulated network. TOB error is measured by RF distance to the true TOB, normalized by the total number of internal edges in the two trees to account for polytomies. ASTRAL guides InPhyNet’s decomposition and provides the starting tree for CAMUS; its restricted subtrees also serve as starting trees for DIMPLE’s MPL subnetwork inference. We measure ASTRAL error as the normalized RF distance to the major displayed tree of the ground-truth network.

##### Network accuracy

To compare network topologies, we use the topological distance of Nakhleh (2010) (Section III, Definition 7), implemented in the `CmpNets` command in PhyloNet (Wen et al. 2018). We normalize this distance by the number of nodes in the ground-truth network.

To complement whole-network topological error, we evaluate hybrid ancestry following the approach of Dai and Molloy (2026). A taxon *x* ∈ *X* is said to have *hybrid ancestry* in a network *N* if there is a directed path in *N* from some reticulation node to the leaf *x*. Let *H*(*N*) and 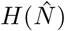 denote the taxa with hybrid ancestry in the true network *N* and the inferred network 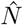, respectively. The *hybrid-ancestry balanced error* (HAE) is defined as

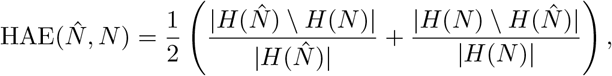

where a term with a zero denominator is defined as zero. Thus, identical hybrid-ancestry sets have error zero, and a tree estimate has error 0.5 when *H*(*N*) ≠ ∅. The measure depends only on which taxa lie below a reticulation, not on the position of the root. HAE consequently does not require rooting at the designated outgroup.

For evaluation using the topological distance of Nakhleh (2010), we rooted each inferred network at the designated outgroup whenever possible. If no valid rooting at the outgroup existed, we used the valid rooting that minimized topological error relative to the ground-truth network. Additional discussion of distance-measure sensitivity is provided in Appendix D.

## 4 Results

### 4.1 Accuracy of gene trees and species phylogenies

Gene-tree estimation error was low overall but higher under high ILS at every dataset size (Table 1). Because the ILS conditions were generated by rescaling network branch lengths, the differences in gene-tree estimation error may reflect the resulting differences in gene-tree branch lengths. TOB error remained low across all conditions, with a mean normalized RF distance below 0.08, and was relatively insensitive to network complexity. In contrast, ASTRAL error increased with network complexity, particularly under high ILS.

**Table 1.** Mean ± standard deviation of normalized RF distances for estimated gene trees, TOBs, and ASTRAL species trees under low and high ILS. Gene-tree errors are summarized by dataset size; TOB and ASTRAL errors are summarized by network level. Gene trees and TOBs are compared with their respective ground truths, whereas ASTRAL trees are compared with the major displayed tree of the ground-truth network. Lower values indicate greater accuracy.

| Estimated input | Condition | Low ILS | High ILS |
| --- | --- | --- | --- |
| Gene trees | 50 taxa | $0.112 \pm 0.016$ | $0.177 \pm 0.027$ |
| | 100 taxa | $0.113 \pm 0.010$ | $0.148 \pm 0.018$ |
| | 250 taxa | $0.120 \pm 0.010$ | $0.143 \pm 0.010$ |
| TOB | Level 1 | $0.026 \pm 0.018$ | $0.042 \pm 0.029$ |
| | Level 2 | $0.030 \pm 0.026$ | $0.052 \pm 0.025$ |
| | Level 3+ | $0.030 \pm 0.020$ | $0.077 \pm 0.050$ |
| ASTRAL | Level 1 | $0.031 \pm 0.022$ | $0.048 \pm 0.031$ |
| | Level 2 | $0.061 \pm 0.037$ | $0.083 \pm 0.039$ |
| | Level 3+ | $0.097 \pm 0.058$ | $0.132 \pm 0.080$ |

### 4.2 Performance of division algorithms

In Experiment 1, **TOB no restriction** preserved all reticulations in nearly all datasets (Fig. 2, left). Under the subset-size constraint, DIMPLE retained all reticulations in 80 of 180 datasets, compared with 31 and 17 for InPhyNet with TINNiK-guided and NJ-based division, respectively. A similar pattern was observed for network level, with DIMPLE showing fewer losses than either InPhyNet division strategy (Fig. 2, right).

**Figure 2.**
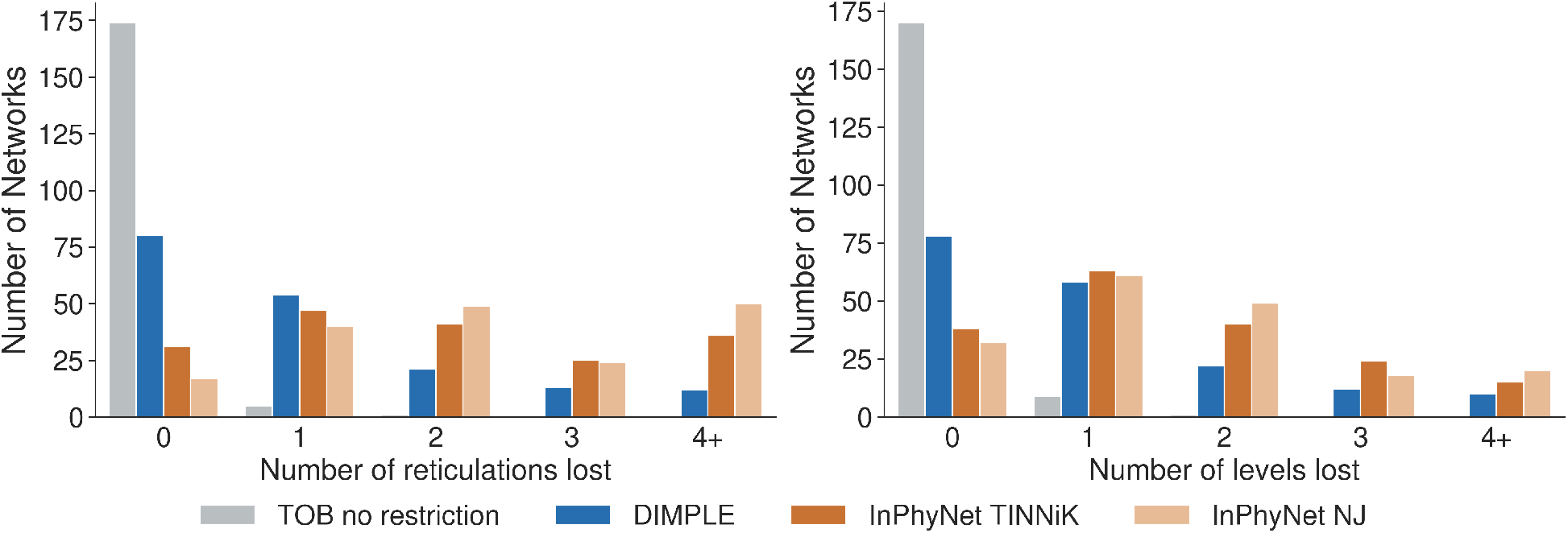
Comparison of division methods in terms of reticulation and level loss across all 180 simulated datasets. Left: the number of datasets with 0 to 4+ reticulations lost. Right: the number of datasets with 0 to 4+ levels lost.

### 4.3 Accuracy of subnetwork inference

To assess subnetwork inference accuracy within each pipeline, we evaluated PhyloNet MPL and SNaQ on the subsets produced by their respective division methods DIMPLE and InPhyNet. The evaluation was restricted to subnetworks overlapping true blobs. Both network inference methods perform comparably (Table S1 in Appendix E). MPL recovers the correct subnetwork in more than half of the cases at levels 1 and 2 (64.5% and 56.9%, respectively) and in 46.6% of cases at level 3+. Error rises with reticulation complexity, with the mean normalized distance increasing from 0.075 at level 1 to 0.138 at level 3+. SNaQ’s exact-recovery rate declines slightly less across levels, and overall its mean normalized error is comparable to MPL’s.

### 4.4 Accuracy of merger on true inputs

With true inputs, the methods differ little on level-1 networks, and the differences grow with network level (Fig. 3). On level-1 networks under low ILS, all four methods achieve a median error of zero under both measures at 50 and 100 taxa. At 50 taxa, DIMPLE exactly recovered 80% of the level-1 networks under low ILS and 90% under high ILS, whereas exact recovery rates for the other methods decreased by 20 to 40 percentage points under high ILS.

**Figure 3.**
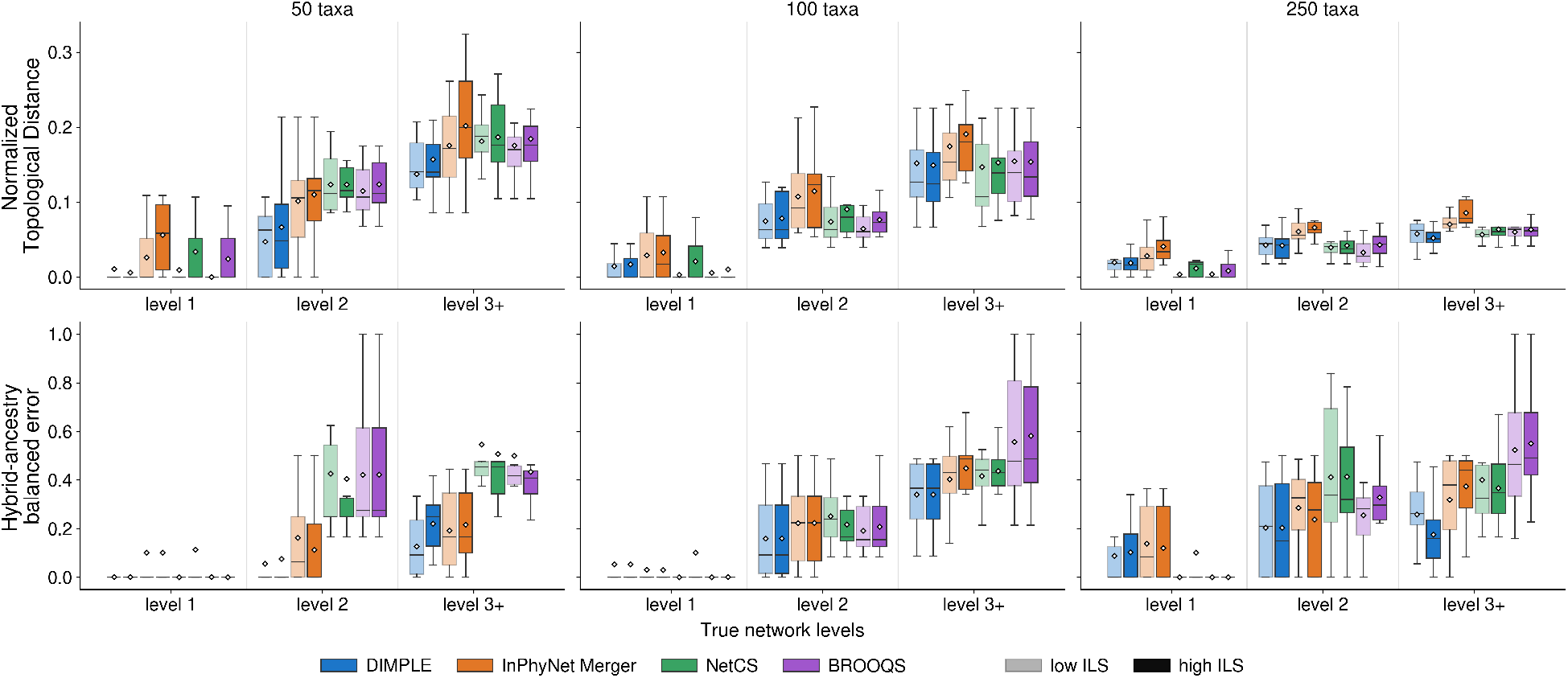
Merger accuracy on true inputs, measured by whole-network normalized topological distance (top row) and HAE (bottom row). DIMPLE and the InPhyNet merger receive the true subnetworks produced by DIMPLE’s division under the true TOB. We include NetCS and BROOQS for comparison; both receive the true TOB and true simulated gene trees. Each panel corresponds to one dataset size, with the three network levels shown along the x-axis. Method identity is encoded by color and ILS setting by opacity: light boxes indicate low ILS, and solid boxes indicate high ILS. Boxes span the interquartile range, the interior line is the median, and the white diamond marks the mean. Each box summarizes 10 replicate networks, except that the InPhyNet merger has 9 at 50 taxa level 3+ (both ILS levels) and at 250 taxa level 2 under high ILS, and NetCS has 9 at 100 taxa level 3+ under low ILS, where those methods produced no output.

DIMPLE maintains high merger accuracy as network complexity increases. Across the 12 higher-level simulation conditions — three dataset sizes, two network levels (2 and 3+), and both ILS levels — it achieves the lowest median HAE in 11, tying with the InPhyNet merger in one of them (Fig. 3, bottom row). This performance is maintained with larger numbers of taxa. For example, on 250-taxon level-3+ networks under high ILS, its median HAE is less than half that of every other method.

Topological distances follow the same ordering with smaller margins: DIMPLE’s median is the lowest or within 0.02 of the lowest median across methods. Its median topological distance is lower than that of the InPhyNet merger in all higher-level conditions. DIMPLE also reconstructs more higher-level networks exactly than other methods, recovering four of the 30 evaluated level-2 networks under low ILS and three under high ILS, compared with one for the InPhyNet merger.

### 4.5 Accuracy of end-to-end network reconstruction

We next compare the complete DIMPLE pipeline with InPhyNet, CAMUS, NetCS, and BROOQS using estimated inputs. ASTRAL serves as a tree-only baseline. For topological error, networks that cannot be rooted at the outgroup (19 for NetCS, 2 for InPhyNet, and 3 for CAMUS) are evaluated using the valid rooting that minimizes their normalized topological distance to the ground-truth network (Section 3.5).

Reconstruction accuracy decreases when estimated inputs are used, but DIMPLE continues to perform well as network complexity increases. On level-1 networks, no method has uniformly lower topological error across all conditions: DIMPLE achieves the lowest median in four of the six level-1 conditions, while at 250 taxa under low ILS, NetCS and BROOQS have lower medians (Fig. 4). On level-1 networks, DIMPLE recovers hybrid ancestry exactly in 60% of networks under low ILS and 53% under high ILS. The corresponding rates decrease from 63% to 33% for NetCS and from 50% to 7% for InPhyNet; BROOQS achieves exact recovery in at most 30% of networks under either ILS condition.

**Figure 4.**
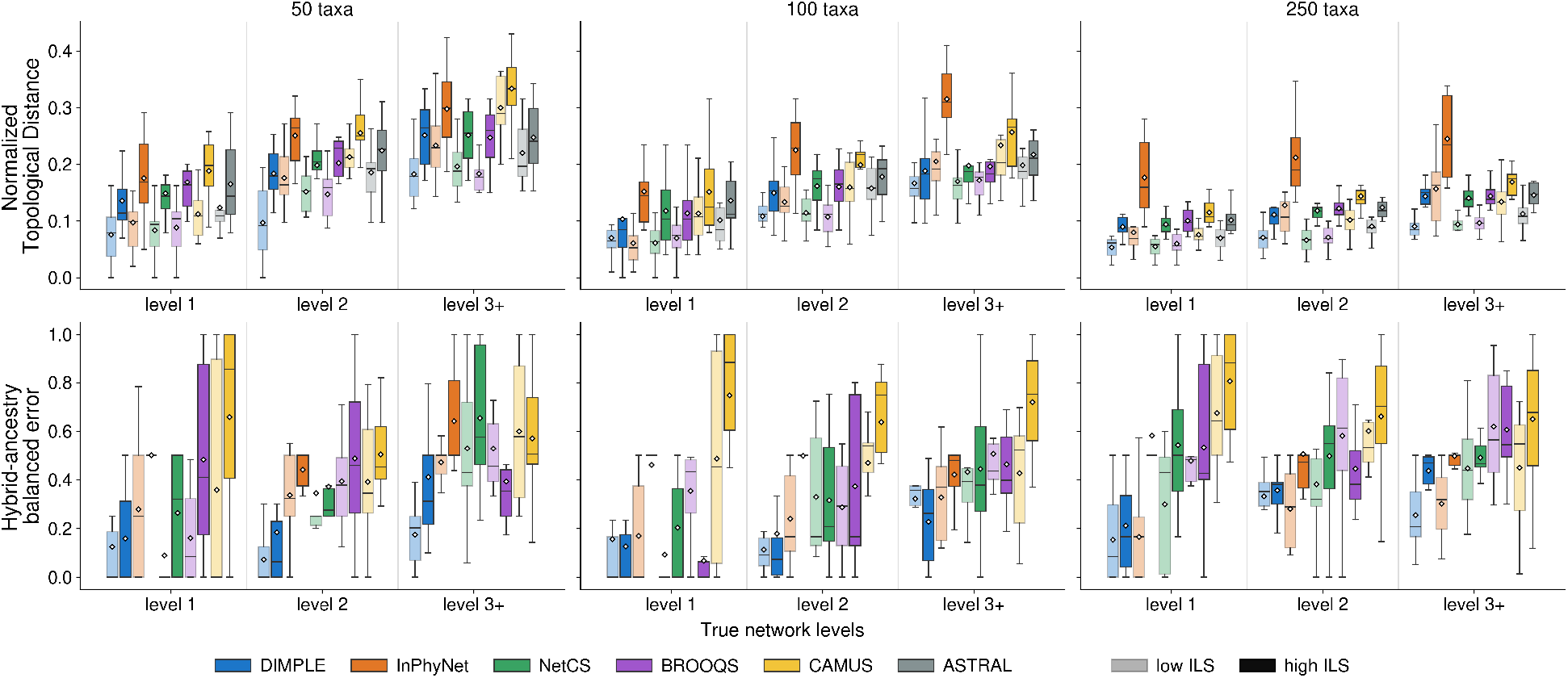
Full-pipeline accuracy of DIMPLE, InPhyNet, NetCS, BROOQS, and CAMUS on estimated inputs. The top row reports normalized topological distance, with the ASTRAL species tree included as a comparison; the bottom row reports HAE, where ASTRAL is omitted because a tree posits no hybrid ancestry and scores 0.5. The maximum subnetwork size is set to 12 for DIMPLE and InPhyNet. Columns correspond to 50, 100, and 250 taxa, and within each panel the boxes are grouped by true network level. Color identifies the method, and opacity identifies the ILS condition: the lighter box of each pair is low ILS and the solid box is high ILS. Boxes span the interquartile range, the interior line is the median, the white diamond marks the mean, and whiskers extend to 1.5 times the interquartile range (outliers not shown). Each box summarizes 10 replicate networks.

On higher-level networks, DIMPLE has lower median topological error than InPhyNet across the evaluated conditions (Fig. 4, top row). When results are pooled across the three dataset sizes, DIMPLE achieves the lowest median HAE on both level-2 and level-3+ networks (Fig. 4, bottom row), recovering hybrid ancestry exactly on 18% of low-ILS and 17% of high-ILS networks, against at most 7% for any other method. Figure S1 compares the numbers of reticulations inferred by DIMPLE and InPhyNet and the number of blobs in the inferred TOB with the true number of reticulations.

### 4.6 CPU time and peak memory usage

We recorded the end-to-end CPU time and peak memory usage for Experiment 3. Runs were performed on a shared computing cluster with 96-core Intel Xeon nodes with 256 GB RAM. InPhyNet ran single-threaded; DIMPLE ran four MPL analyses in parallel with four threads each (16 threads in total), and CAMUS used its automatic parallelization with 16 threads.

At 50 taxa, NetCS and BROOQS (4.4 min each) and CAMUS (6.4 min) need less than a tenth of DIMPLE’s CPU time (78.5 min) and InPhyNet’s (112.7 min) (Fig. 5). The methods scale very differently. From 50 to 250 taxa, CAMUS’s CPU time increases nearly 1,500-fold to 9,344.7 min, 25 times DIMPLE’s. NetCS and BROOQS grow about 30-fold (136.7 min and 135.1 min); both totals are dominated by TOB-QMC (131.8 min at 250 taxa), which is why they are nearly identical. DIMPLE grows only 5-fold (369.7 min) and at 250 taxa is substantially faster than InPhyNet (663.0 min). At 250 taxa, TOB-QMC accounts for about a third of DIMPLE’s total CPU time. The CPU time spent on MPL is essentially flat from 100 to 250 taxa: 209.5 min at 100 taxa and 223.9 min at 250 taxa. DIMPLE’s cost increases sharply with network level: at 50 taxa, it increases from 16.5 min at level 1 to 240.0 min at level 3+.

**Figure 5.**
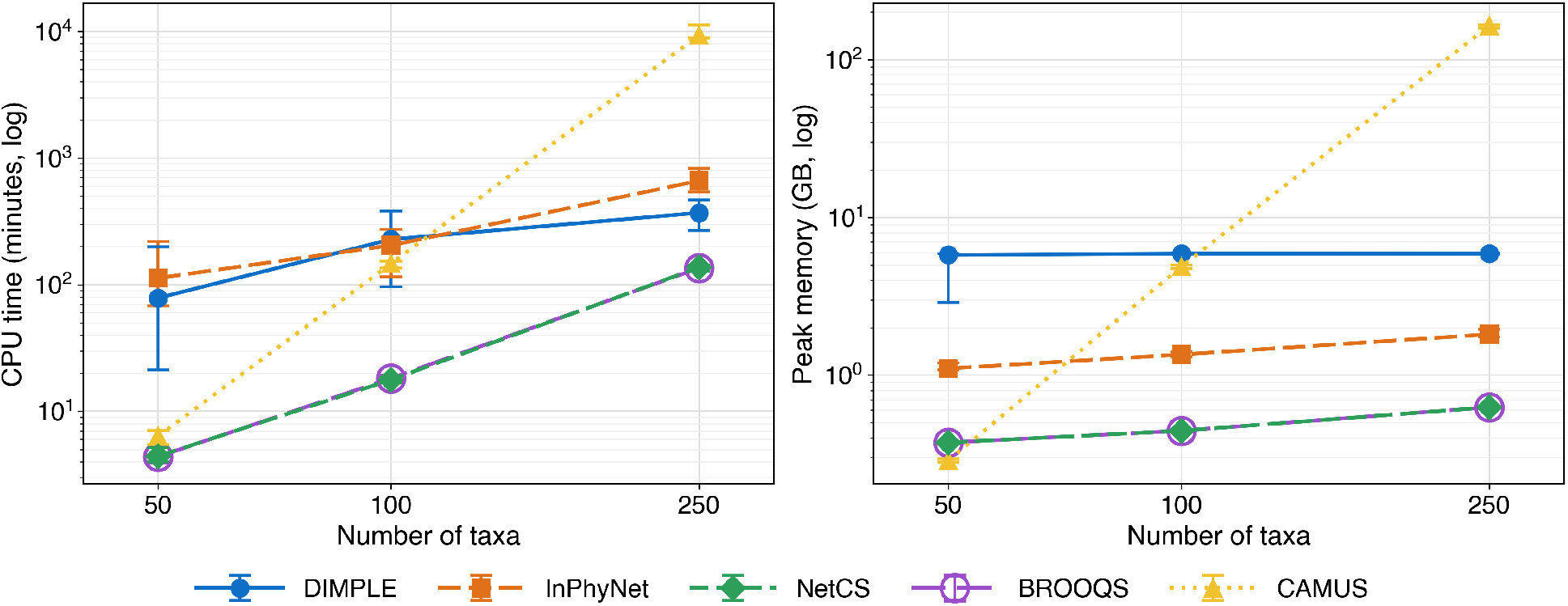
Computational cost of DIMPLE, InPhyNet, NetCS, BROOQS, and CAMUS as the number of taxa grows on all simulated data. Points represent medians, and error bars span the 25th to 75th percentiles. Left: end-to-end CPU time (minutes). It includes building the ASTRAL species tree and the TOB for methods that require them. Right: peak memory, measured as the maximum resident set size (RSS). BROOQS and NetCS share TOB-QMC, which sets their peak, so their points coincide (BROOQS is drawn as an open circle).

BROOQS and NetCS use the least memory at 100 and 250 taxa, staying below 0.7 GB, while InPhyNet grows from 1.3 to 1.8 GB. DIMPLE’s peak is nearly constant at about 6 GB across different numbers of taxa, reflecting four MPL analyses running in parallel (about 1.5 GB each). CAMUS’s peak memory usage increases from 297 MB at 50 taxa, the lowest of all methods, to 164 GB at 250.

### 4.7 Results on biological data

The biological datasets and inference settings are described in Sections 3.2 and 3.3. Fig. 6 shows the networks inferred by DIMPLE. Networks inferred by NetCS, BROOQS, CAMUS, and InPhyNet are shown in Figs. S7, S8, S9, and S10, respectively, with further comparisons in Appendix I. CPU time and peak memory usage on the biological datasets are compared in Fig. S11.

**Figure 6.**
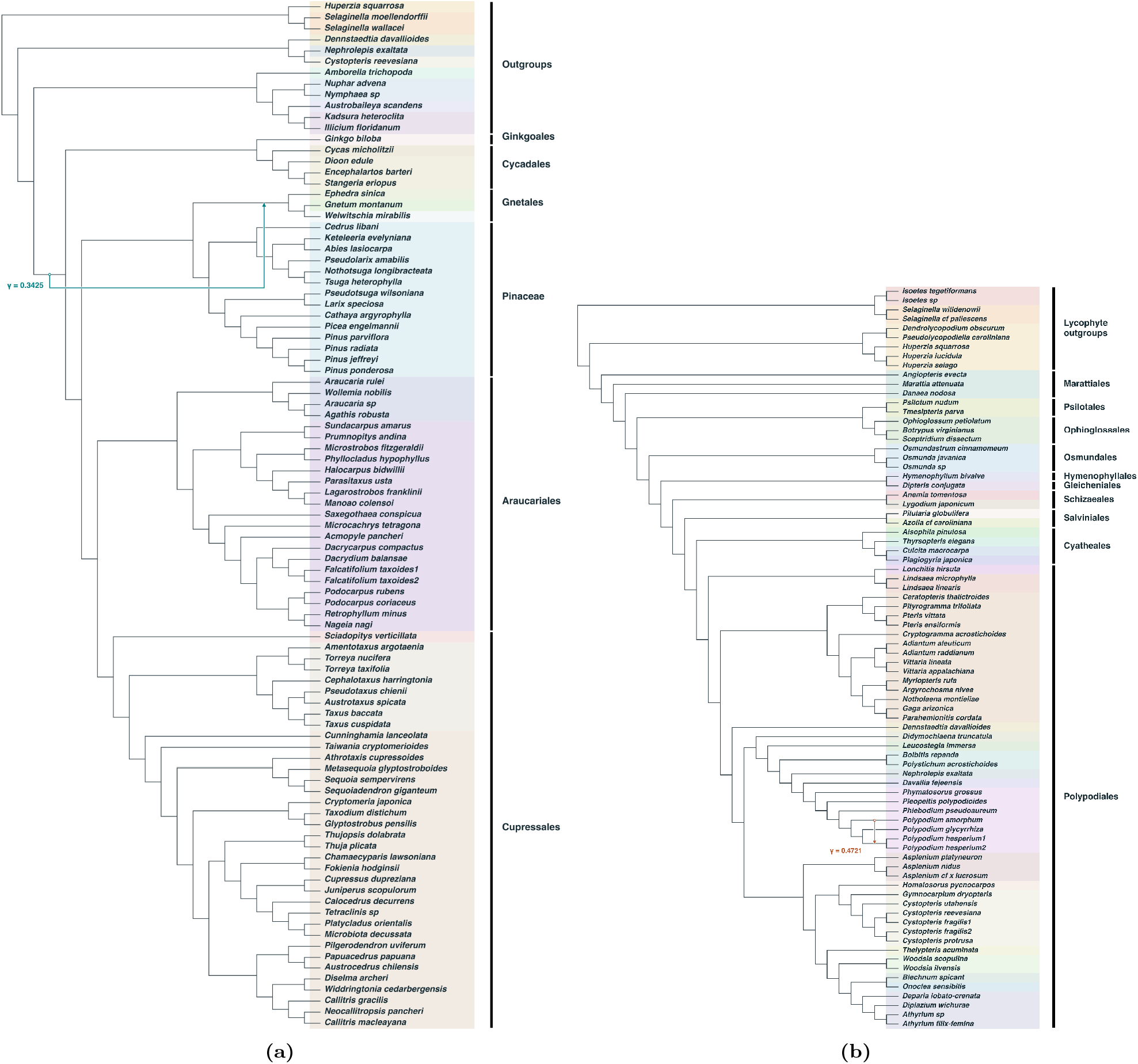
Phylogenetic networks inferred by DIMPLE on the gymnosperm and fern subsets of the 1KP transcriptome dataset. Colors group taxa by family, and black bars indicate orders. Colored arrows indicate the reticulation events labeled with their inheritance probabilities *γ*. (a) Gymnosperm. Gnetales (*Gnetum, Welwitschia, Ephedra*) is recovered as a hybrid lineage with two parental connections: one places Gnetales sister to Pinaceae, while the other places it sister to the remaining gymnosperms, representing conflicting placements of Gnetales in the inferred network. NetCS recovers the same network topology. (b) Fern. One reticulation is recovered within *Polypodium*.

Across the two subsets, DIMPLE recovers two reticulations spanning distinct biological scenarios: one in the fern subnetwork and one in the gymnosperm subnetwork.

#### Polypodium

The two *Polypodium hesperium* tips in the fern network descend from a hybrid node with parental edges to *P. amorphum* and *P. glycyrrhiza*. This matches one of the well-documented allopolyploidy events in ferns: *P. hesperium* is an allotetraploid derived from these two diploid progenitors. Plastid and nuclear sequence data support this origin and indicate that hybridization occurred in both directions between the parental species (Sigel et al. 2014).

#### Gnetales

The evolutionary placement of Gnetales (*Gnetum, Welwitschia, Ephedra*) remains one of the longest-standing controversies in seed plant phylogeny. Three major hypotheses have been proposed: the Gnetifer hypothesis places Gnetales as sister to all conifers; the Gnepine hypothesis places them as sister to Pinaceae; and the Gnecup hypothesis places them as sister to the remaining conifers excluding Pinaceae. Different genomic datasets have supported different hypotheses, suggesting substantial conflict among loci rather than a single unambiguous evolutionary history (Bowe et al. 2000; Ran et al. 2018; Stull et al. 2021). DIMPLE reconstructs Gnetales as a reticulate lineage with two parental connections: one places Gnetales sister to Pinaceae, consistent with the Gnepine hypothesis, while the other places it sister to the remaining gymnosperms, outside the conifer crown. Rather than selecting a single hypothesis, the inferred network captures the conflicting phylogenetic signal present across gene trees, but this result alone does not confirm a reticulate origin of Gnetales.

## 5 Discussion

In this work, we introduced DIMPLE, a divide-and-conquer framework for scalable phylogenetic network inference. DIMPLE uses an inferred TOB to guide taxon set decomposition, applies MPL to infer subnetworks, and merges them into a network on the full taxon set. The decomposition step partitions the set of taxa into subsets according to the TOB structure and repeats this process to increase reticulation coverage. Our merger combines subnetworks on overlapping taxon sets by constructing a tree backbone and adding reticulations supported by the subnetworks when they sufficiently improve pseudo-likelihood. Together, these components enable inference without an explicit limit on the level of the final network.

Our simulation results support using the TOB to guide the division step for supernetwork inference. Reticulation can complicate the gene-tree signal used to estimate a single species tree with methods such as ASTRAL or NJ. In our experiments, the inferred TOB remained accurate as network complexity increased, whereas the ASTRAL tree diverged further from the major displayed tree of the ground-truth network. Using a fully resolved species tree to guide the division step in our pipeline could lead to reticulation loss across taxon partitions. While all methods with explicit subset-size caps can lose some reticulation signal, DIMPLE retained the most reticulations among methods with the same limit.

With the true TOB and true subnetworks, DIMPLE reconstructs level-1 networks nearly perfectly and maintains high merger accuracy on higher-level networks. The comparison with the InPhyNet merger highlights the effectiveness of DIMPLE’s merging strategy, including its ability to combine overlapping subnetworks: both receive true subnetworks and simulated gene trees, yet DIMPLE achieves lower topological and hybrid-ancestry errors on higher-level networks. NetCS (Dai and Molloy 2026) and BROOQS (Arasti and Mirarab 2026) also achieve high accuracy on level-1 networks when given the true TOB, consistent with their reported results, but their hybrid-ancestry accuracy is lower on higher-level networks. A higher-level blob contains multiple reticulations, so representing it as a single cycle cannot preserve all of its reticulation structure. DIMPLE’s ability to infer multiple reticulations within a blob contributes to more accurate reconstruction of hybrid ancestry.

For methods evaluated with both true and estimated inputs, reconstruction accuracy generally decreases when estimated inputs are used. DIMPLE maintains low hybrid-ancestry error on higher-level networks and achieves lower median topological error than InPhyNet across the evaluated higher-level conditions. Together, these findings suggest that resolving individual blobs with NetCS and BROOQS is effective for level-1 networks when the TOB is accurate, whereas DIMPLE’s subnetwork inference and merging offer advantages for recovering hybrid ancestry on higher-level networks with either true or estimated inputs. The comparison between true and estimated inputs also highlights the importance of inference accuracy.

Additional studies of maximum subnetwork size (Appendix H) show that increasing the maximum subset size from 6 to 12 improves reconstruction accuracy, whereas increasing it further to 18 provides limited and inconsistent additional benefits. We therefore use 12 as a practical compromise between retaining reticulation information and keeping subnetwork inference manageable. MPL subnetwork inference is a major contributor to DIMPLE’s runtime, making the size and complexity of individual subproblems important determinants of its computational requirements. Empirically, the CPU time of the MPL step rises sharply with network level and increases less than proportionally across the evaluated dataset sizes. At 250 taxa, DIMPLE requires roughly 6 CPU hours, little more than half of InPhyNet’s time.

Our biological analyses demonstrate the scalability of DIMPLE on phylogenomic datasets and its ability to recover biologically meaningful reticulate relationships. In ferns, DIMPLE recovers the parental relationships underlying the established allopolyploid origin of *Polypodium hesperium*, linking it to *P. amorphum* and *P. glycyrrhiza*. In gymnosperms, rather than forcing a single bifurcating placement for Gnetales, the inferred network represents the longstanding conflict among competing evolutionary hypotheses.

Inference beyond level-1 networks is difficult, and the theoretical limits of divide-and-conquer network inference remain unresolved. In particular, here we present a heuristic that performs well on the examined data. However, a follow-up study into what classes of level-*k* supernetworks are identifiable in divide-and-conquer frameworks is warranted. For divide-and-conquer, the question remains whether accurate subnetworks on the selected taxon subsets, together with the guide and other auxiliary information, contain sufficient information to determine the full network. Further work should characterize the classes of networks and ideal data inputs for which such reconstruction is identifiable.

Our current ways of quantifying network inference accuracy are limited. Current measures of topological accuracy of inferred phylogenetic networks can obscure errors in the rare reticulation events when tree-like structure dominates the network. They can be sensitive to minor tree inconsistencies: a single subtree-prune-and-regraft (SPR) move can produce a large Robinson–Foulds distance. In our study, ASTRAL, a species *tree* estimation method, achieves a median topological error of 0.128 for low ILS, comparable to those of network inference methods. Thus, low topological error alone does not establish accurate reticulation recovery. We therefore use hybrid-ancestry error as a complementary assessment, and we plan to develop more robust measures of network similarity.

Several directions could improve divide-and-conquer inference. Joint inference of the backbone and reticulation structure could improve accuracy, although implementing such an approach would be challenging and would require carefully defined graph operations. Allowing the merger to revise erroneous subnetwork relationships can also improve robustness. Moreover, a limitation of our simulation evaluation is that we supply the true number of reticulations for subnetwork inference; future work is needed to investigate automatic selection of reticulation counts and statistical support for inferred reticulations. Together, these tasks could form part of an agentic iterative workflow: estimate the TOB, divide the set of taxa, infer subnetworks, merge them, and reassess the resulting network to determine whether further division or inference is needed. Developing such a pipeline rather than a single algorithmic tool would provide a systematic approach to refining network estimates through repeated inference and re-evaluation. Broader evaluation across dataset sizes, ILS levels, and simulation conditions would help assess its effectiveness.

## 6 Data Availability

The simulated networks, biological datasets, and outputs from all evaluated methods will be available on Zenodo. DIMPLE is open source at https://github.com/NakhlehLab/DIMPLE.

## 7 Acknowledgments

This work was supported in part by the National Science Foundation grants DMS/NIGMS-2153704 and DBI-2030604.

## 7.1 AI Assistance

We used Claude (Anthropic) and Codex (OpenAI) to assist with coding and editing manuscript text. The authors reviewed and revised all AI-assisted content and take responsibility for the final manuscript.

## A Method availability

DIMPLE is open source at https://github.com/NakhlehLab/DIMPLE. The workflow uses TOB-QMC (Dai et al. 2026), implemented in TREE-QMC v4.0.1 (Han and Molloy 2023), PhyloNet v3.8.2 (Wen et al. 2018), and ASTRAL v5.7.8 (Zhang et al. 2018).

The example commands below use `gene_trees.tre`, containing one Newick gene tree per line on the full taxon set, and OUT as the outgroup label. The file `tob_rooted.tre` denotes the inferred TOB rooted at the designated outgroup before division. These examples illustrate the workflow; the settings used in each experiment are described in Section 3.3.

### Tree of blobs (TOB)

#### Step 1: Infer a base tree and annotate its branches

First, we estimate the base tree from the gene trees, apply the quartet hypothesis tests around each branch, and store the resulting minimum *p*-value on that branch. The default iteration limit for the *p*-value search is two times the number of taxa squared, while the authors of TREE-QMC recommend setting the iteration limit to one fourth the number of taxa squared for large data sets.

~~~
tree-qmc --iter_limit_blob 5000 --store_pvalue \
-i gene_trees.tre -o base_tree_pvalues.tre
~~~

#### Step 2: Construct and root the TOB

Contract the annotated branches using the stored *p*-values with the default thresholds *α* = 10^−7^ and *β* = 0.95, and root the result at the outgroup:

~~~
tree-qmc --blob --alpha 1e-7 --beta 0.95 --load_pvalue \
--root OUT -i base_tree_pvalues.tre -o tob_rooted.tre
~~~

### ASTRAL

ASTRAL provides the starting tree for subnetwork inference:

~~~
java -jar astral.5.7.8.jar \
-i gene_trees.tre -o astral.tre -t 4
~~~

### DIMPLE

The DIMPLE workflow consists of division, subnetwork inference with PhyloNet, and merging. All three stages share one output directory chosen by the user, named `dimple_out` in the commands below.

### Division

~~~
python -m dimple.divider.generate_k_divisions \
--tob tob_rooted.tre --output_dir dimple_out \
--size 12 --k 5 --seed 0 --outgroup OUT
~~~

#### Subnetwork inference

Every subnetwork is inferred individually. `--list` prints each division run with the indices of its subsets. A subnetwork can then be inferred for each subset with the number of reticulations chosen for it; the inference results for each tested reticulation limit are retained. Finally, `--assemble` writes the division run’s `subnets.txt` from the results, taking for each subset the network inferred with the listed number of reticulations (one number per subset in order).

*List the subsets in each division run*.

~~~
python -m dimple.phylonet.infer_subnetworks dimple_out \
--gene-trees gene_trees.tre --base-tree astral.tre \
--phylonet PhyloNet.jar --list
*Infer a subnetwork for one subset at a specified reticulation count*.
python -m dimple.phylonet.infer_subnetworks dimple_out \
--gene-trees gene_trees.tre --base-tree astral.tre \
--phylonet PhyloNet.jar --pl 4 \
--only blob00/run_000 --subset 1 --max-ret 1
*Assemble the selected subnetworks for a division run*.
python -m dimple.phylonet.infer_subnetworks dimple_out \
--gene-trees gene_trees.tre --base-tree astral.tre \
--phylonet PhyloNet.jar \
--only blob00/run_000 --assemble 1,1,0
~~~

Omitting `--only` and `--subset` runs subnetwork inference for every subset using the same `--max-ret` value.

#### Merging

Run the merger using the assembled subnetworks:

~~~
python -m dimple.merger.merger_full_pip dimple_out \
--tob tob_rooted.tre --gene-trees gene_trees.tre \
--max-runs 10 --out-name full_merger
~~~

The final network is written to `dimple_out/full_merger/merged_full.nwk`.

These example settings use a maximum subset size of 12 taxa, five division attempts per megablob, and random seed 0. For subnetwork inference, the user specifies the maximum number of reticulations allowed; the example sets this limit to 1. For a more thorough analysis, users can repeat inference with limits of 0, 1, 2, 3, or higher and examine how log pseudo-likelihood improves as additional reticulations are allowed. These score profiles can inform manual selection of the reticulation count. Each search uses the ASTRAL tree restricted to the subset taxa as its starting tree.

## B Algorithm details

Throughout this section, *T*_*N*_ denotes the input tree of blobs, *b* denotes a megablob of *T*_*N*_, and *G* denotes the input gene trees. The collection *S* contains the taxon subsets used for subnetwork inference within megablobs, and *P* contains the pruned subtrees.

### B.1 Division algorithms

#### Algorithm S1

Divide (*T*_*N*_, *s, K*)

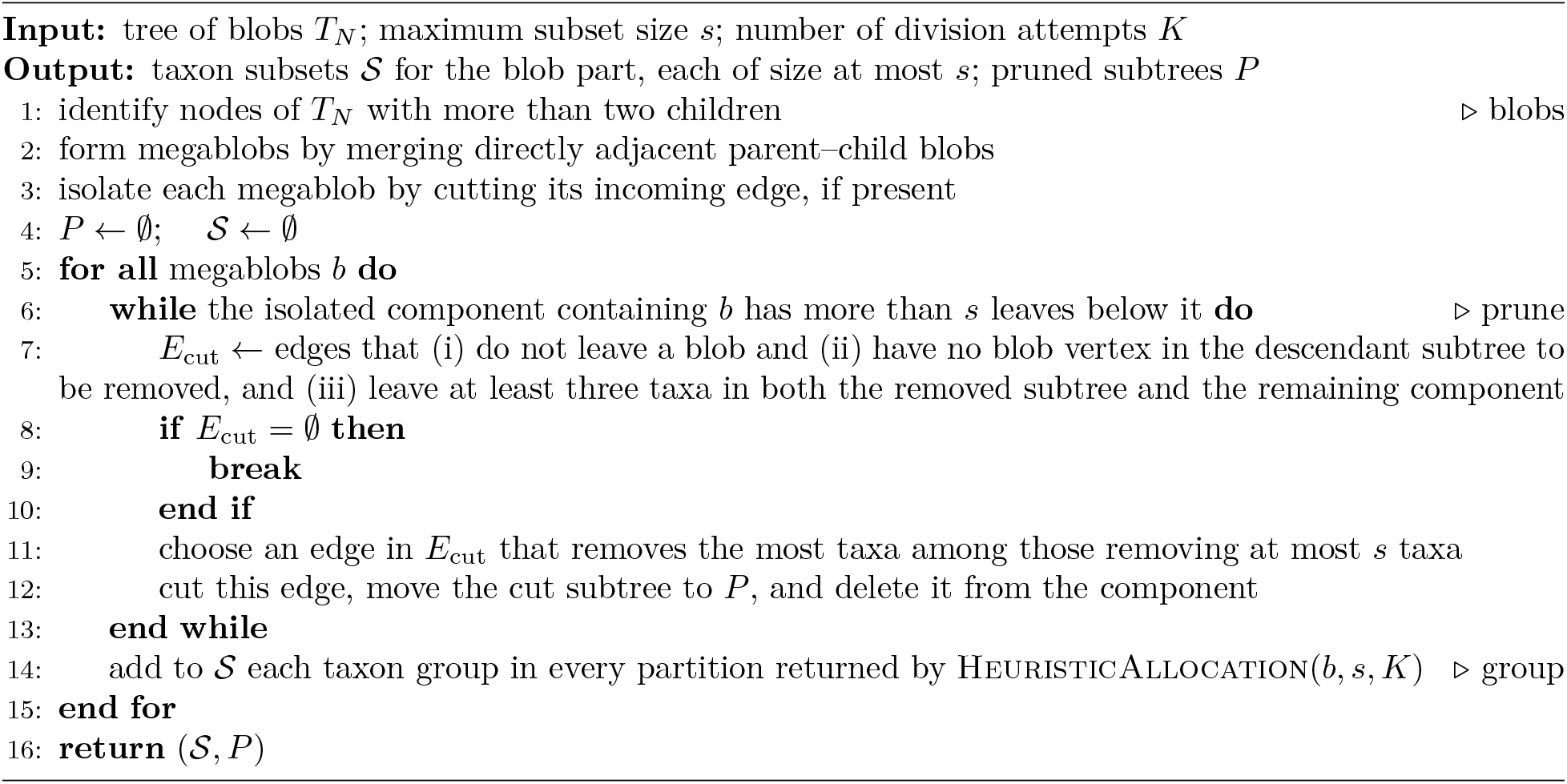

Let *X*_*b*_ denote the set of taxa remaining in the isolated component of megablob *b* after pruning.

#### Algorithm S2

HeuristicAllocation (*b, s, K*)

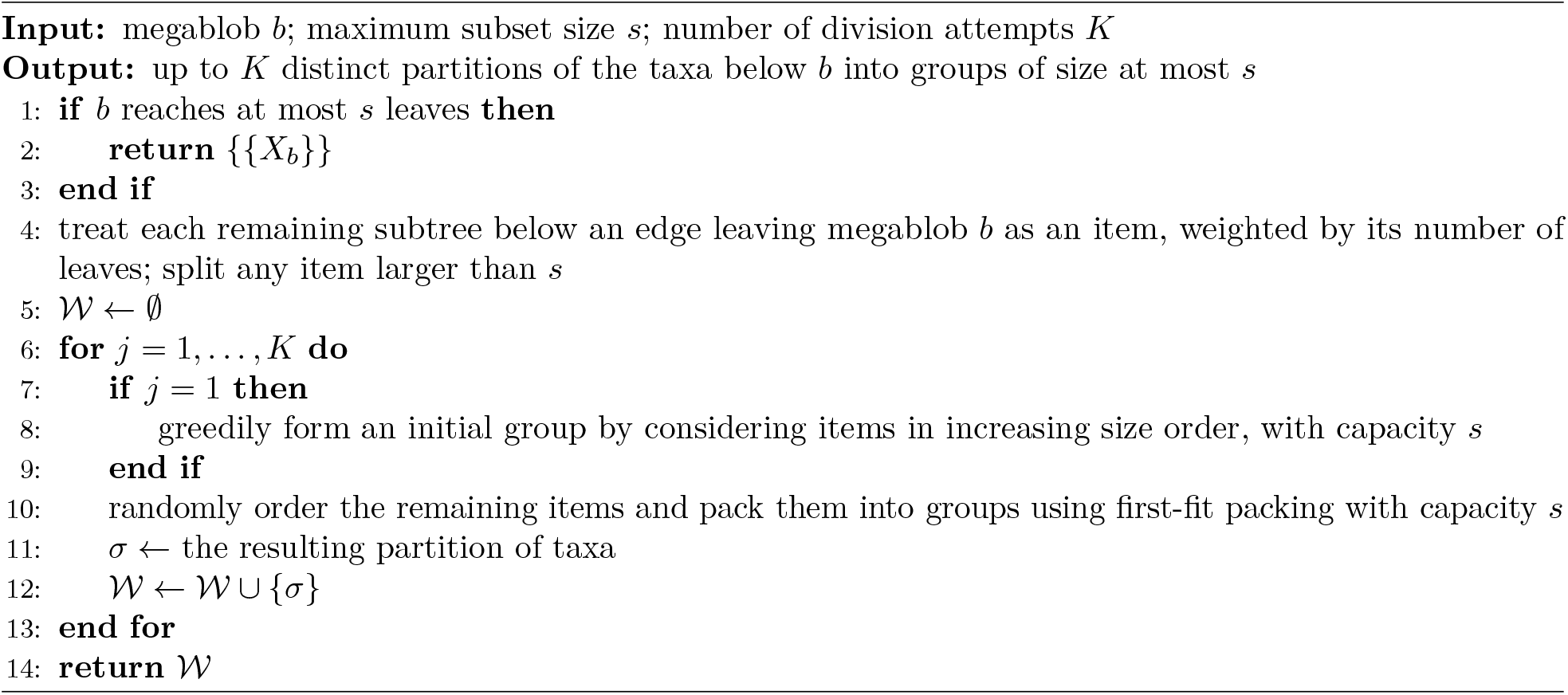

### B.2 Merger algorithms

Let *N* denote the collection of all inferred subnetworks, and let N_*b*_ denote those associated with megablob *b*.

#### Algorithm S3

Merge (*T*_*N*_, *N*, *G, P*)

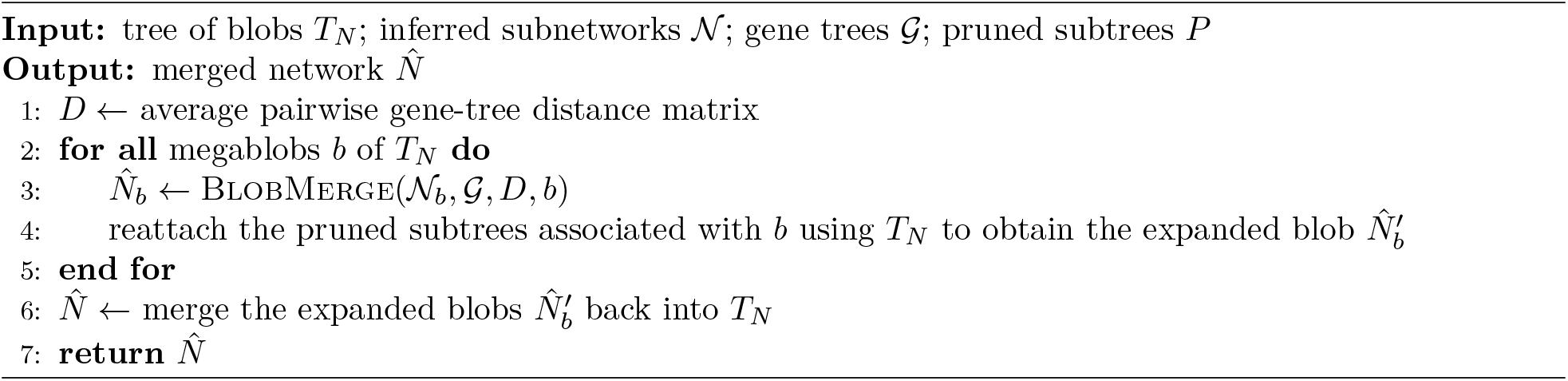

#### Algorithm S4

Blobmerge (*N*_*b*_, *G, D, b*)

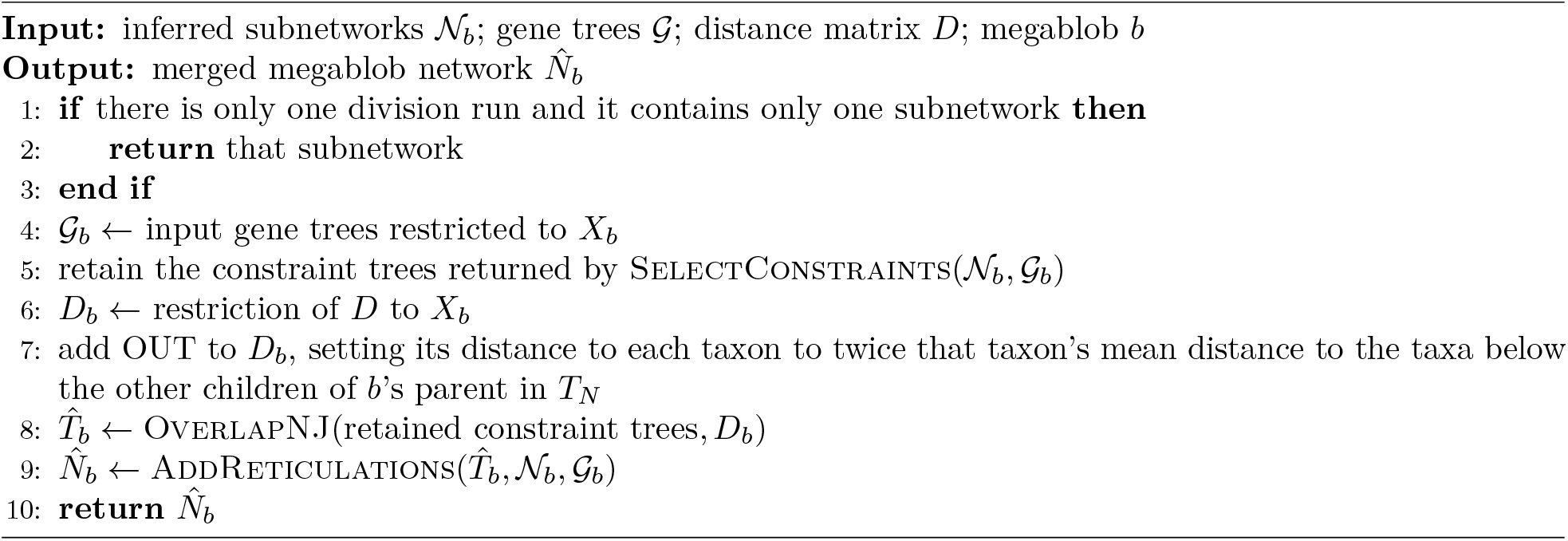

#### Algorithm S5

SelectConstraints (*N*_*b*_, *G*_*b*_)

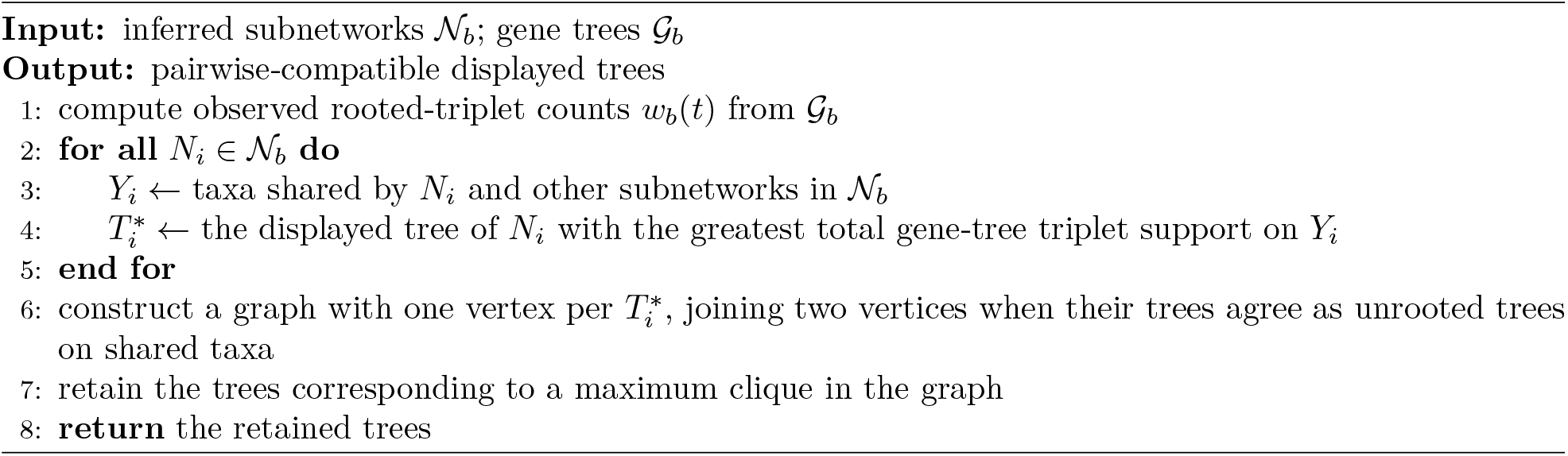

For *m* current trees, define the neighbor-joining criterion (Studier and Keppler 1988)

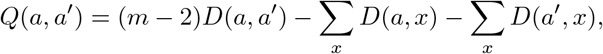

where *D*(*a, a*^′^) is the current distance between the trees represented by labels *a* and *a*^′^, and both sums range over all current tree labels. Pairs with smaller *Q* are considered first.

We use the unrooted topologies of the selected displayed trees for the OverlapNJ step. Before neighbor joining, we add a temporary outgroup taxon OUT to the distance matrix. For each taxon *x* ∈ *X*_*b*_, we set its distance to OUT to twice the mean of *D*(*x, y*) over the external sibling taxa *y* of megablob *b*, which are the taxa below the other children of *b*’s parent in *T*_*N*_. The constraint trees do not contain OUT, so its placement is guided by its distances without imposing an additional topological constraint on its position. After neighbor joining, we root the resulting tree on the edge incident to OUT and then remove OUT to obtain the rooted backbone 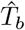 on *X*_*b*_ before the next step.

#### Algorithm S6

OverlapNJ(constraint trees, *D*)

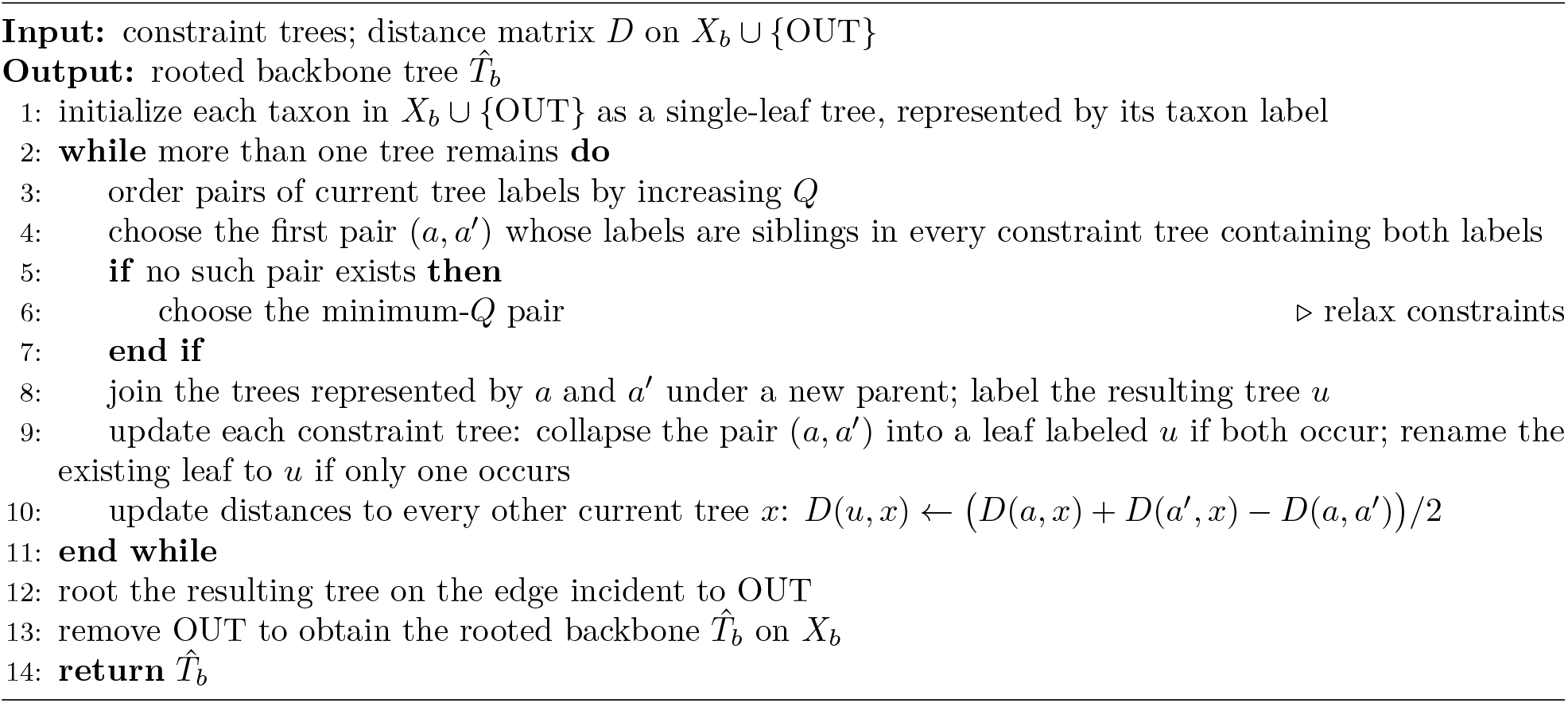

For reticulation insertion, let *f* (*N*; *G*_*b*_) denote the negative log pseudo-likelihood of network *N* evaluated using the gene trees *G*_*b*_; smaller values indicate a better fit. Because *G*_*b*_ remains fixed, we abbreviate *f* (*N*; *G*_*b*_) as *f* (*N*).

For each reticulation, we identify the taxa descended from the other child of each of its two parents. We group reticulations when these taxon sets overlap for both corresponding parents. For each group, we select the subnetwork of the reticulation with the largest descendant-taxon set. All reticulations from the selected subnetworks are then considered for insertion. For each candidate, we identify which parental relationship is already represented in the backbone. We refer to the other parent in its source subnetwork as the parent to be inserted. Inserting a reticulation can change which taxa descend from an edge and therefore affect the attachment locations available to later candidates. Here, a candidate is considered before another if its descendant taxa overlap the taxa descended from the other child of the second candidate’s parent to be inserted.

We evaluate candidate networks using the rooted-triplet pseudo-likelihood implementation in PhyNetPy, a Python library for phylogenetic network analysis (PhyNetPy). Because backbone construction does not estimate branch lengths, we initially assigned all branches a length of 1. Preliminary experiments with branch-length optimization yielded only small improvements in log pseudo-likelihood while increasing runtime. We therefore kept branch lengths fixed at 1 during merger scoring; inheritance probabilities were copied from the source subnetworks and held fixed.

For each candidate or already accepted reticulation *r*, let *X*(*r*) denote the set of taxa descended from the corresponding reticulation node in its source subnetwork. Let *N* be the current merged network, and let *d*_*N*_ (*u, v*) denote shortest-path distance in its underlying undirected graph. For a candidate *r*, let *v* be the child endpoint of its best valid attachment edge in *N*. We classify *r* as *nearly redundant* if there exists an already accepted reticulation node *r*^′^ such that *X*(*r*) ∩ *X*(*r*^′^) ≠ ∅ and *d*_*N*_ (*v, r*^′^) ≤ 2.

For nearly redundant candidates, the best placement must decrease the negative log pseudo-likelihood by at least 2% relative to the current score (*τ* = 0.02). All other candidates are accepted whenever their best placement strictly decreases the score (*τ* = 0).

#### Algorithm S7

AddReticulations 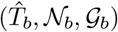

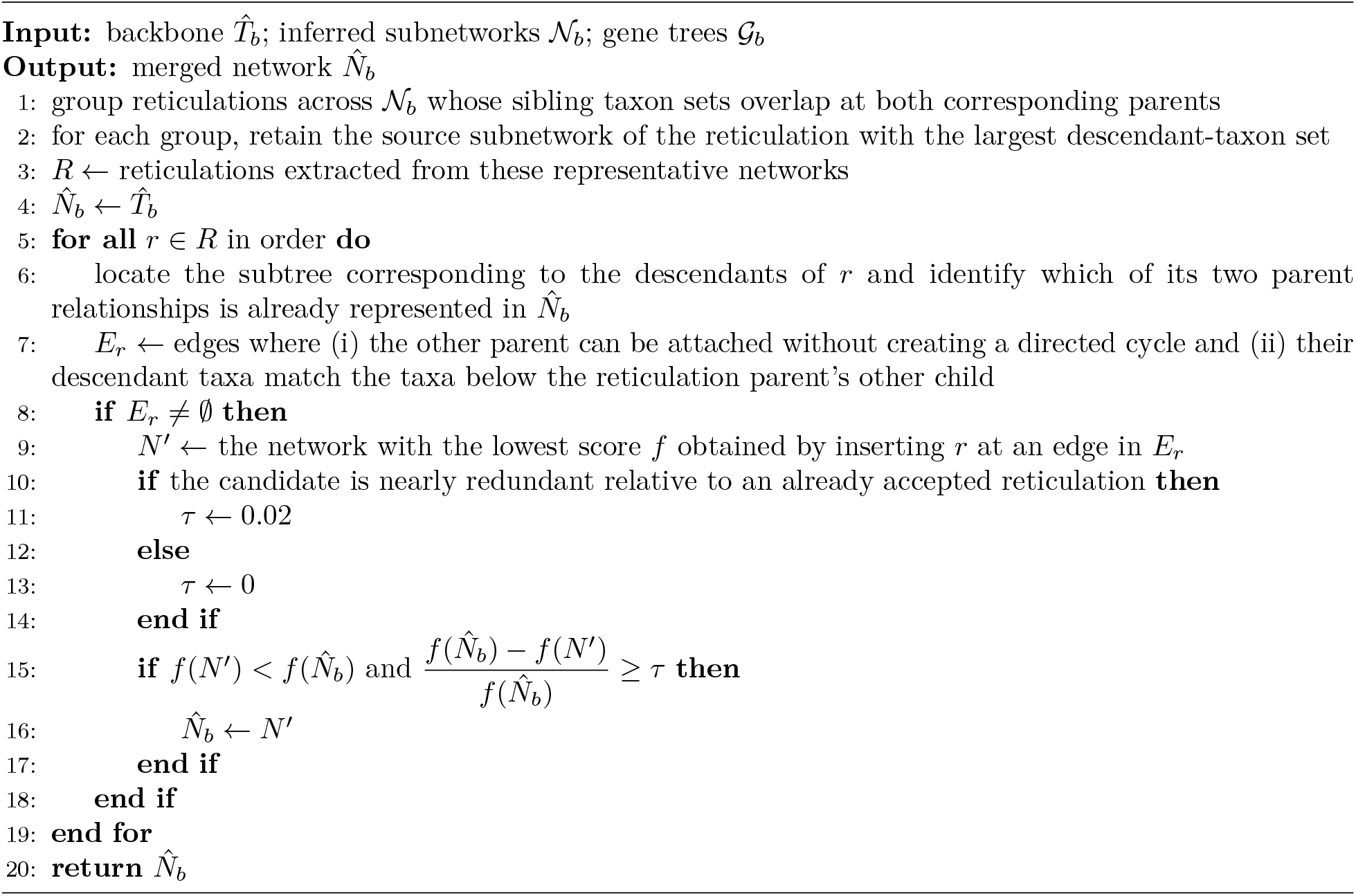

## C Runtime complexity

Let *n* = |*X*| be the number of taxa, *q* the number of gene trees, *s* the maximum subset size, and *K* the number of division attempts. For each megablob *b*, let *n*_*b*_ = |*X*_*b*_|, let *m*_*b*_ be the number of input subnetworks, and let *k*_*b*_ ≤ *m*_*b*_ be the number of retained constraint trees. The following analysis covers the main division and merger operations described in Algorithms S1–S7 in the appendix.

### TOB inference

We infer the TOB using TOB-QMC (Dai et al. 2026). Dai et al. report a runtime of *O*(*qn*^3^) under the assumptions of their analysis, where *n* is the number of taxa and *q* is the number of input gene trees.

### Division

Identifying blob vertices requires a traversal of the TOB and takes *O*(*n*) time. The complete division procedure is not a single traversal as the algorithm repeatedly searches for an eligible subtree to prune. With leaf counts and blob membership computed by a postorder traversal, each candidate-edge sweep takes linear time. There can be *O*(*n*) cuts, giving a worst-case bound of *O*(*n*^2^) for these repeated sweeps.

For grouping, let *a*_*b*_ be the number of items for megablob *b* after splitting any item larger than *s*. Sorting the items takes *O*(*a*_*b*_ log *a*_*b*_) time. Each first-fit attempt processes *a*_*b*_ items and may scan up to *a*_*b*_ existing groups per item, giving 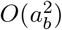 worst-case time. The *K* packing attempts therefore cost 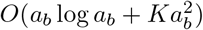 per megablob.

### Distance matrix and displayed-tree selection

The average pairwise gene-tree distance matrix is computed once for the full dataset in *O*(*q* · *n*^2^) time. Algorithm S5 additionally computes rooted-triplet support and enumerates displayed trees. There are 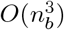 possible rooted triplets on *X*_*b*_. Their counts can be computed in 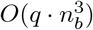 time. These counts are computed once per megablob and reused when scoring its candidate displayed trees.

If input subnetwork *N*_*i*_ has *h*_*i*_ binary reticulations, our displayed-tree enumeration considers up to 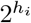 parent-edge choices. Each resulting tree is then scored on its shared taxa. Scoring each candidate requires extracting triplets from a tree with at most *s* taxa and evaluating them, costing 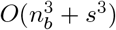.

### Compatible-subset filtering

The compatibility graph has *m*_*b*_ vertices and requires 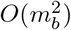 pairwise comparisons, each between trees with at most *s* taxa. The subsequent maximum-clique search has exponential worst-case complexity in *m*_*b*_. In practice, both *m*_*b*_ and *s* are small, so compatibility filtering and maximum-clique selection are fast per megablob.

### Overlapping neighbor joining

Algorithm S6 performs *O*(*n*_*b*_) joins. At a stage with *m* current trees, computing and sorting the *O*(*m*^2^) candidate pairs costs *O*(*m*^2^ log *m*). Checking each pair against up to *k*_*b*_ constraint trees may take *O*(*k*_*b*_) time per pair. Including distance and constraint updates, the joining stage is bounded by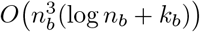. This reduces to 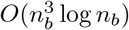 when the number of constraint trees is treated as fixed.

### Reticulation insertion

Let *r*_*b*_ be the number of candidate reticulations processed by Algorithm S7. A binary network on *n*_*b*_ taxa with at most *r*_*b*_ reticulations has *O*(*n*_*b*_ + *r*_*b*_) edges, so each candidate may need to be evaluated at *O*(*n*_*b*_ + *r*_*b*_) insertion locations in the worst case. Let *C*_*b*_ denote an upper bound on the time required to evaluate the pseudo-likelihood of a single candidate network for megablob *b*, after gene-tree triplet frequencies have been precomputed. Scoring the candidate placements therefore requires *O*(*r*_*b*_(*n*_*b*_ + *r*_*b*_)*C*_*b*_) time.

## D Sensitivity of network distance measures

Comparing two phylogenetic networks is harder than comparing two trees. A single misplaced reticulation edge can change the set of displayed trees, the cluster system, and the rooted-subnetwork structure at the same time. Different dissimilarity measures respond to these changes in very different ways. One commonly used measure is the *cluster distance*, defined as the size of the symmetric difference between the hardwired (or softwired) cluster sets of two networks. However, cluster distance is only a pseudo-metric on network topologies: it can be zero for non-isomorphic networks. As a simple example, consider the two level-1 networks

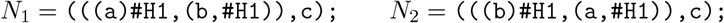

which differ only in the hybrid child. Let Cl(*N*) denote the cluster set of *N*. A direct calculation shows that both induce the same cluster set,

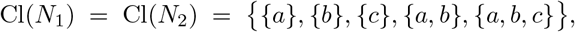

under both the hardwired and softwired definitions. Thus the cluster distance satisfies *d*_Cl_(*N*_1_, *N*_2_) = 0, even though the topological difference is obvious. The distinct asymmetry between *N*_1_ and *N*_2_ is lost because the same leaf clusters are present in both networks no matter which leaf is the hybrid child.

We chose the topological distance metric over cluster-based alternatives because cluster distance does not clearly reflect missing reticulations. A displayed tree can share many clusters with its source network despite containing no reticulations, so a low cluster distance need not imply accurate reticulation recovery. By contrast, the topological distance metric distinguishes nodes whose children differ between the two networks, so removing a reticulation changes the relevant equivalence classes and directly increases the score.

## E Subnetwork inference error

**Table S1:**
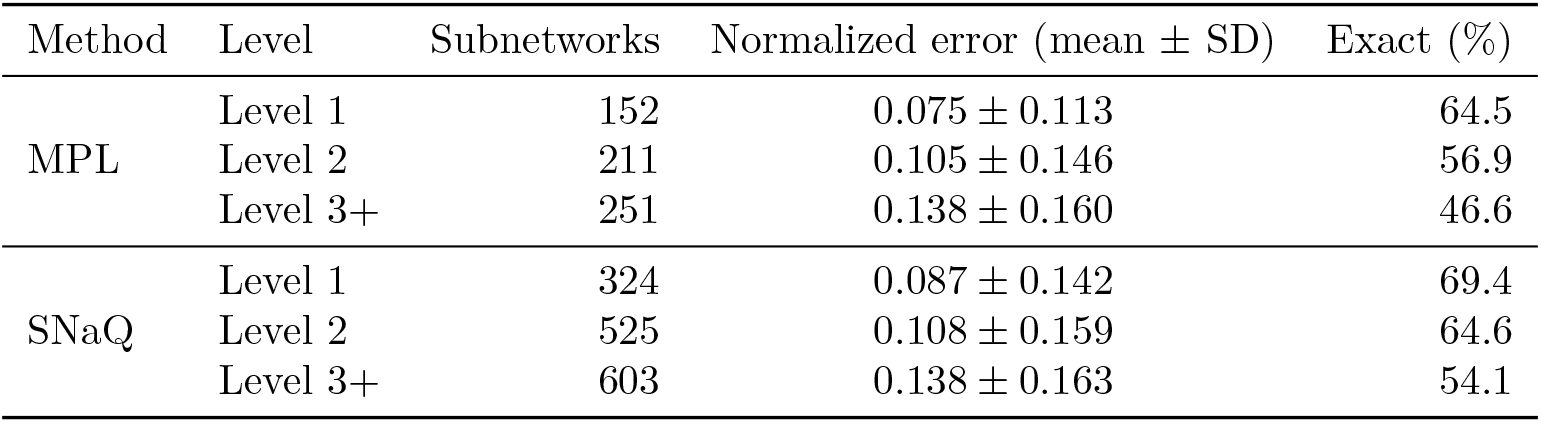
Subnetwork inference error by level of the ground-truth network for MPL (on DIMPLE’s division) and for SNaQ (on InPhyNet’s division, restricted to subnetworks overlapping true blobs). This is not a controlled comparison because inference uses the respective division schemes and therefore different taxon subsets. Error is the topological distance between the inferred and true subnetworks, normalized by the number of nodes in the true subnetwork. SNaQ estimates unrooted networks, so each SNaQ subnetwork is evaluated at the valid rooting that minimizes its normalized topological distance to the true subnetwork.

## F Number of inferred reticulations

We compared three quantities: the number of reticulations inferred by DIMPLE, the number of reticulations inferred by InPhyNet, and the number of blobs in the inferred TOB (Fig. S1). CAMUS was excluded from this comparison because we selected its output network based on the true number of reticulations. BROOQS infers one reticulation per inferred blob, whereas NetCS does so only for blobs of size greater than four, leaving smaller blobs unresolved.

Across the 180 inferred networks with estimated gene trees, DIMPLE and InPhyNet generally underestimated the number of reticulations. DIMPLE’s median matched the true number for networks with one or two reticulations but fell below it for networks with three or more. DIMPLE recovered the number of reticulations exactly in 35% of replicates. InPhyNet consistently had lower medians than DIMPLE. The median number of inferred blobs was at or above the true number of reticulations for networks with one to three reticulations but fell below it for networks with four or more. However, a blob in a higher-level network can contain multiple reticulations, so even an accurately inferred TOB need not have as many blobs as the network has reticulations.

## G Effect of the number of division runs

### G.1 Division algorithm performance

As *K* increases, more reticulations are recovered and the gap to TOB-no-restriction narrows. Adding division runs retains more reticulations: DIMPLE-5, DIMPLE-10, and DIMPLE-15 retain all reticulations on 80, 96, and 105 networks, steadily narrowing the gap to the unconstrained TOB-no-restriction reference (Fig. S2).

**Figure S1:**
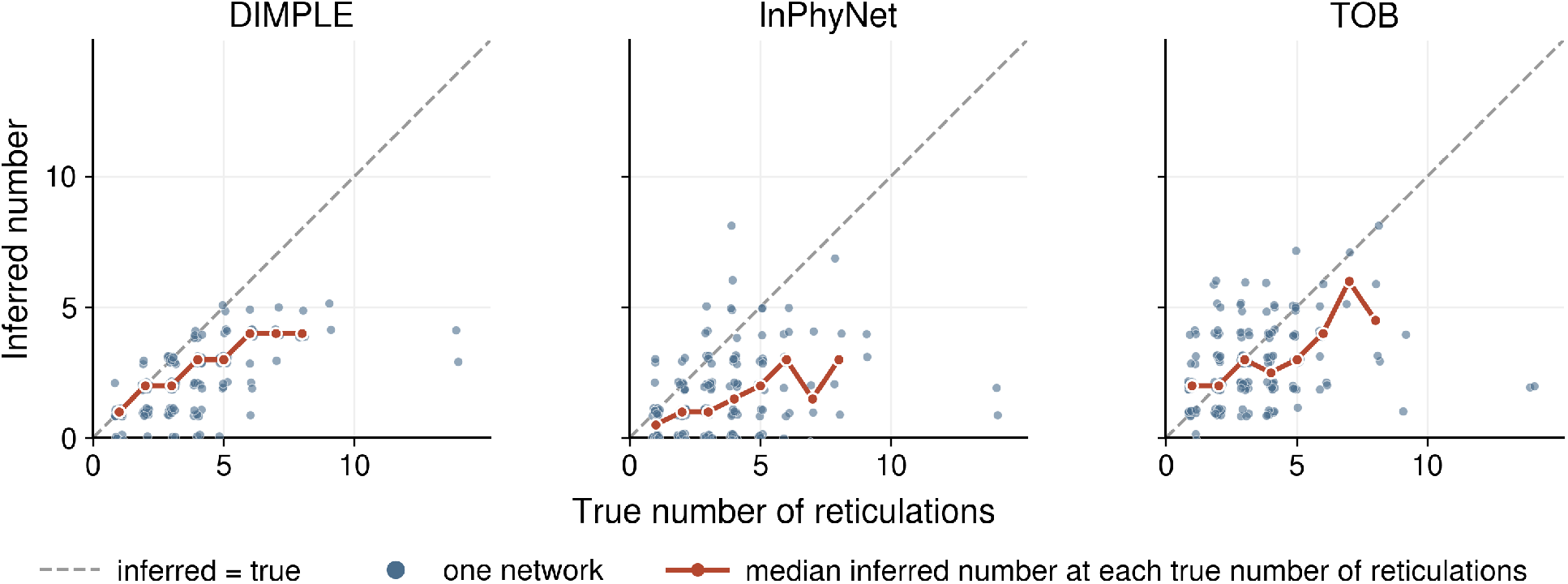
Numbers of reticulations inferred by DIMPLE and InPhyNet and numbers of blobs in the inferred TOBs, plotted against the true number of reticulations across 180 replicates inferred with estimated gene trees. Each point represents one replicate network. Solid lines with circular markers connect the median values. The dashed diagonal marks equality with the true number of reticulations.

**Figure S2:**
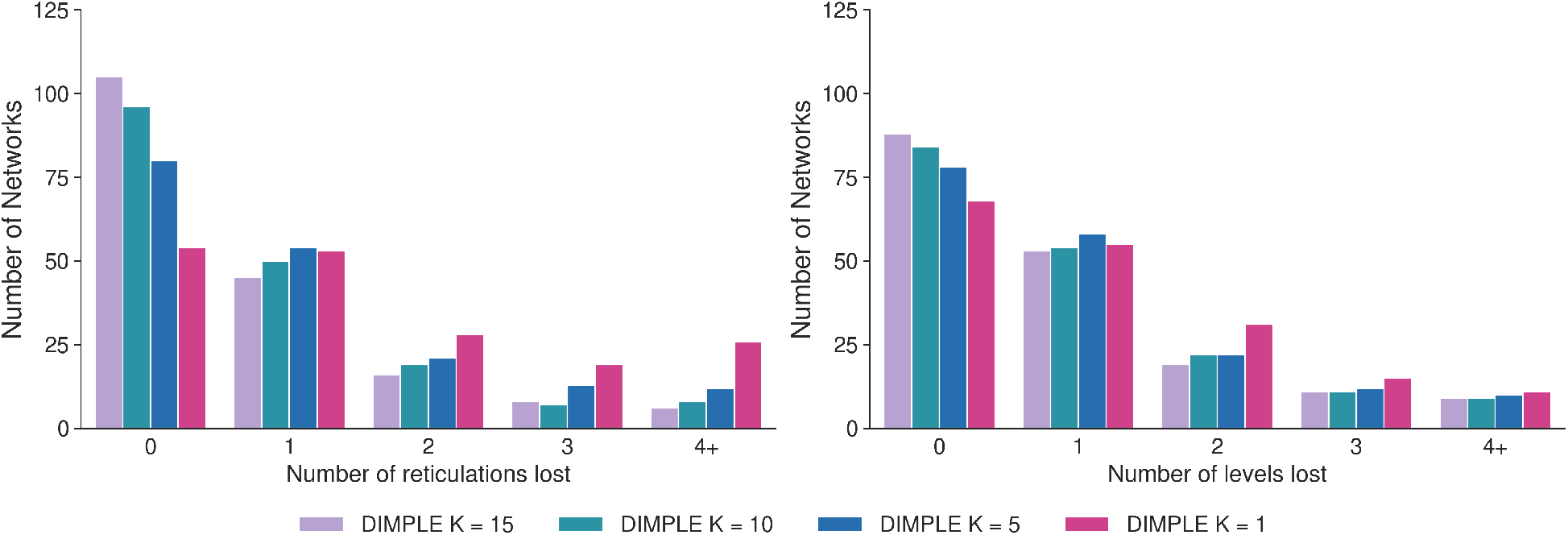
Effect of the number of division runs *K* on reticulation retention during division. Left: reticulations lost. Right: decrease in network level. The bars show counts across all 180 simulated datasets.

### G.2 Merger accuracy with true and estimated inputs

Under true inputs, the pooled median HAE falls from 0.26 at *K* = 1 to 0.05 at *K* = 5 and 0.00 at *K* = 10 under low ILS, and from 0.25 to 0.07 and 0.00 under high ILS. On level-1 networks the median HAE is zero at every evaluated value of *K*, except at 250 taxa under low ILS, where it falls from 0.08 at *K* = 1 to 0.00 at *K* = 5. The largest gains with increasing *K* are on level-3+ networks. Averaged across dataset sizes and ILS conditions, the decrease in median HAE between *K* = 1 and *K* = 10 is 0.29 on level-3+ networks, compared with 0.15 on level-2 networks.

Under estimated inputs, larger *K* reduces error in most higher-level settings. On 100-taxon level-2 networks under low ILS, the median HAE falls from 0.13 at *K* = 1 to 0.00 at *K* = 10.

**Figure S3:**
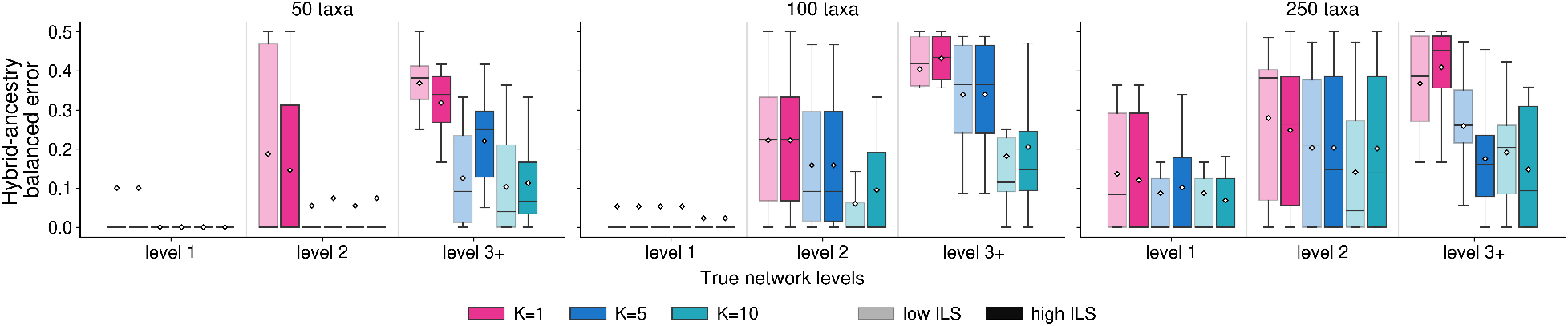
Effect of the number of division runs *K* on DIMPLE accuracy given true subnetworks, using hybrid-ancestry balanced error between the inferred and ground-truth networks. Columns correspond to dataset sizes and the x axis to the level of the true network. Color indicates *K*; within each color, the lighter box shows low ILS and the darker box high ILS. Boxes span the interquartile range, the interior line marks the median, and the white diamond marks the mean. Each box summarizes 10 replicate networks.

**Figure S4:**
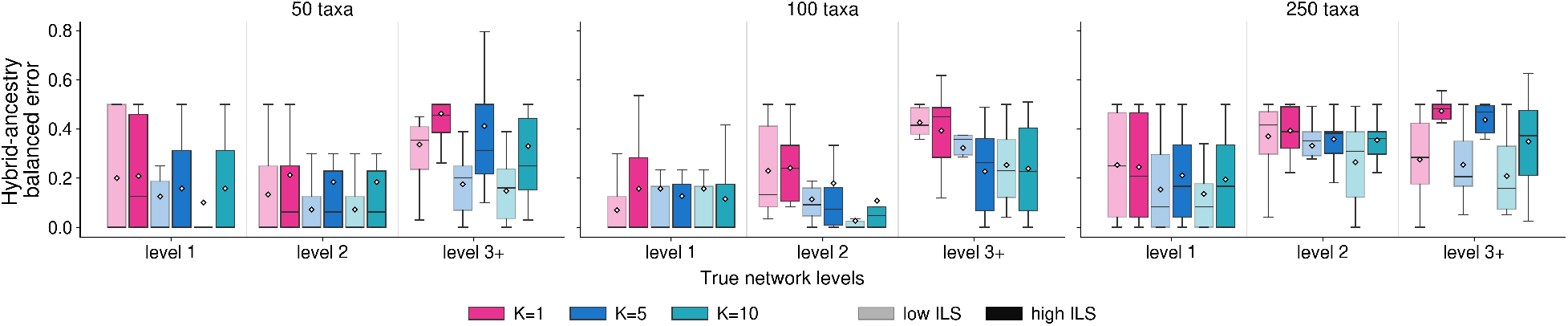
Effect of the number of division runs *K* on end-to-end accuracy, using hybrid-ancestry balanced error between the inferred and ground-truth networks. Columns correspond to dataset sizes and the x axis to the level of the true network. Color indicates *K*; within each color, the lighter box shows low ILS and the darker box high ILS. Boxes span the interquartile range, the interior line marks the median, and the white diamond marks the mean. Each box summarizes 10 replicate networks.

**Figure S5:**
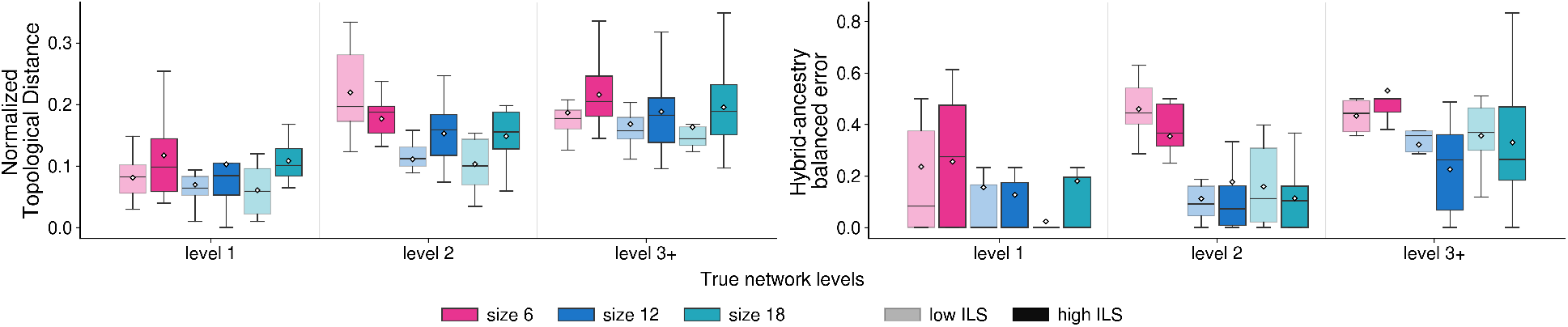
Normalized topological error (left) and hybrid-ancestry balanced error (right) of DIMPLE on 100-taxon networks as the maximum division size takes values of 6, 12, and 18. Color indicates the maximum subset size; within each color, the lighter box shows low ILS and the darker box high ILS. Boxes span the interquartile range, the interior line marks the median, and the white diamond marks the mean. Each box summarizes 10 replicate networks.

## H Effect of maximum subnetwork size on accuracy

Increasing the maximum subset size from 6 to 12 improved DIMPLE’s end-to-end accuracy at every network level and under both ILS conditions (Fig. S5). The gain was largest for hybrid ancestry: on level-2 networks, the median balanced error fell from 0.45 (low ILS) and 0.37 (high ILS) at size 6 to 0.09 and 0.07 at size 12, while the median topological distance fell from 0.20 to 0.11 under low ILS and from 0.19 to 0.16 under high ILS. Beyond 12, the benefit was small: size 18 slightly lowered the median topological distance under low ILS at every level, but slightly raised it under high ILS at levels 1 and 3+, and it did not reduce the median hybrid-ancestry error. We therefore chose 12 as the default maximum subset size, capturing the substantial improvement over size 6 while keeping subnetwork inference computationally manageable.

## I Supplementary results on the 1KP dataset

### I.1 Reticulation-count selection

For each fern and gymnosperm subnetwork, we examined the highest log pseudo-likelihood attained as the number of reticulations increased. Larger gains indicate a greater improvement in fit, while smaller gains indicate diminishing improvement. We used the size of these gains to choose the number of reticulations, starting from a tree and adding one reticulation at a time. Where the first reticulation gave only a negligible improvement, we reported the tree; otherwise we continued adding reticulations until the improvement became small. We marked the selected number of reticulations with a gray vertical line in each plot.

### I.2 Comparison with other methods

On the fern and gymnosperm subsets of the 1KP transcriptome dataset (One Thousand Plant Transcriptomes Initiative 2019), we compared DIMPLE with InPhyNet, CAMUS, NetCS, and BROOQS. For InPhyNet, we used the same taxon subsets recorded in InPhyNet’s supplementary information; we ran SNaQ inference at a single reticulation value per subset following the reticulation count reported in the original paper. For CAMUS, we followed the authors’ recommendation to select the number of reticulations by examining where the optimization score stops improving substantially. We took the one-reticulation network: the first hybrid branch accounts for 42.4% (fern) and 64.1% (gymnosperm) of satisfied quartets, while each subsequent branch contributes under 7%.

Both datasets are challenging for every method: each has fewer than 400 gene trees, and polyploidy in plants further complicates reticulation inference.

#### Gymnosperm

DIMPLE, NetCS, and BROOQS recover a Gnetales reticulation connecting a Pinaceae-side parent with a deeper gymnosperm lineage. BROOQS identifies three additional hybrid nodes as it fully resolves the TOB. The InPhyNet output has two reticulations: one involving Gnetales and one within Cupressaceae. CAMUS instead connects the *Stangeria* lineage to Araucariales and Cupressales.

**Figure S6:**
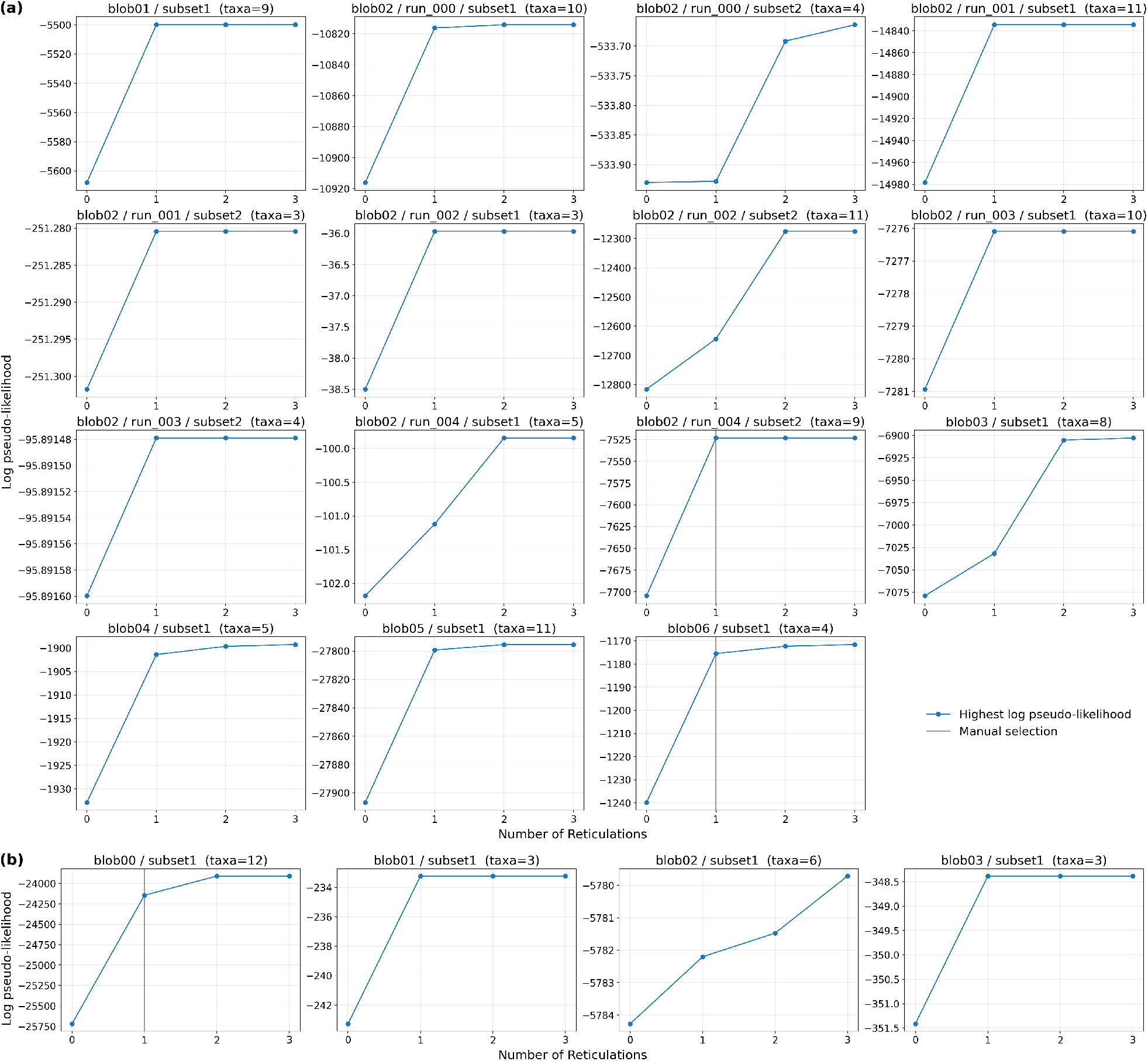
Selecting the number of reticulations per subnetwork on (a) the fern dataset and (b) the gymnosperm dataset. Each panel is one subnetwork, labeled by its megablob, leaf subset, and the number of taxa. The solid curve with point markers shows the highest log pseudo-likelihood attained by any network with at most that number of reticulations, across all `PhyloNet InferNetwork_MPL` searches (hmax = 0, …, 3). The gray vertical line marks the number of reticulations selected for subnetwork inference.

**Figure S7:**
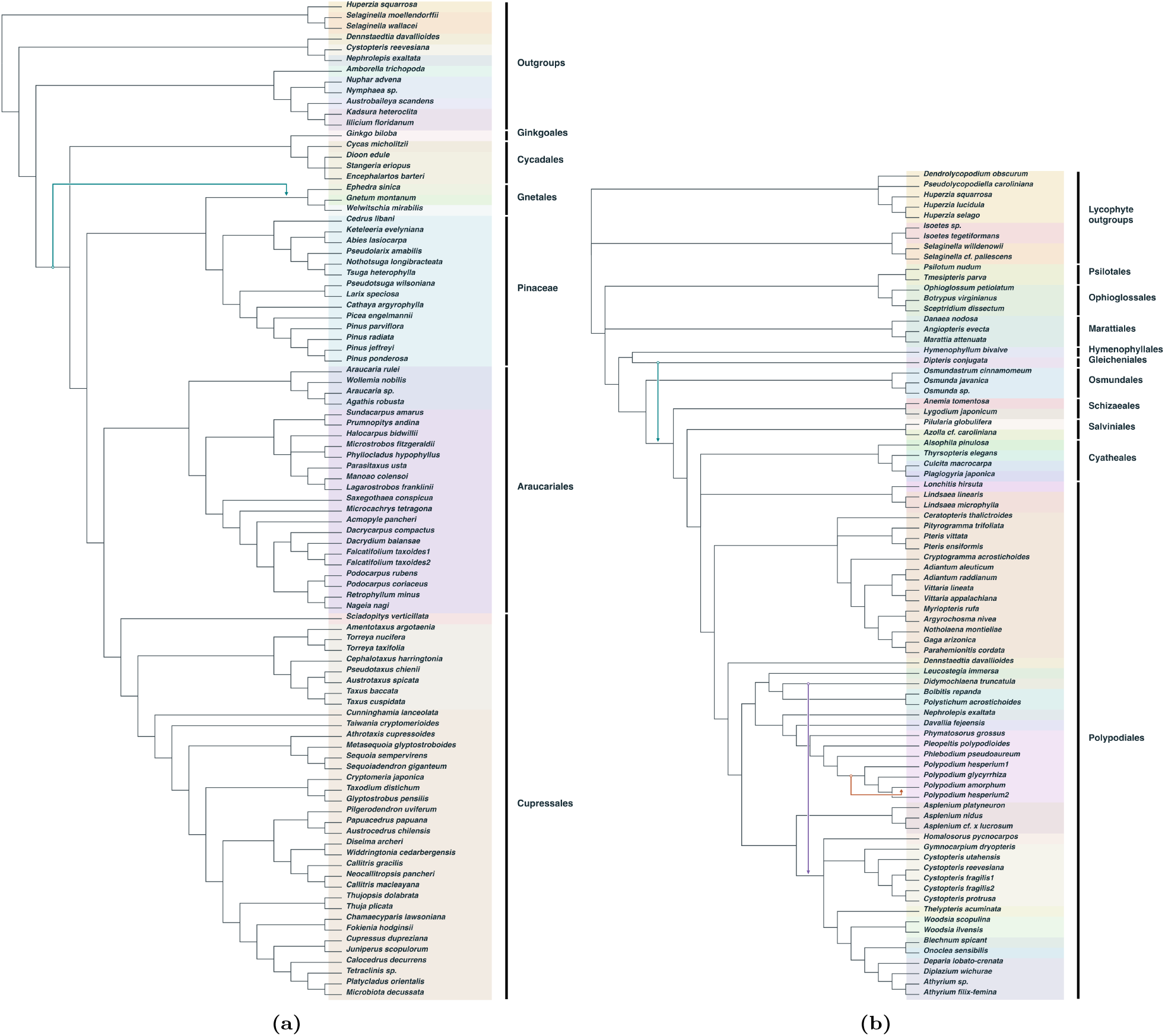
Networks inferred by NetCS on the gymnosperm and fern subsets of the 1KP dataset. (a) Gymnosperm subset. (b) Fern subset.

#### Fern

DIMPLE, InPhyNet, NetCS, and BROOQS recover a reticulation in the *Polypodium* region, but identify different hybrid nodes: *P. hesperium* in DIMPLE, *Phlebodium pseudoaureum* in InPhyNet, and *P. amorphum* in NetCS and BROOQS. CAMUS instead recovers a reticulation spanning the core leptosporangiate ferns.

**Figure S8:**
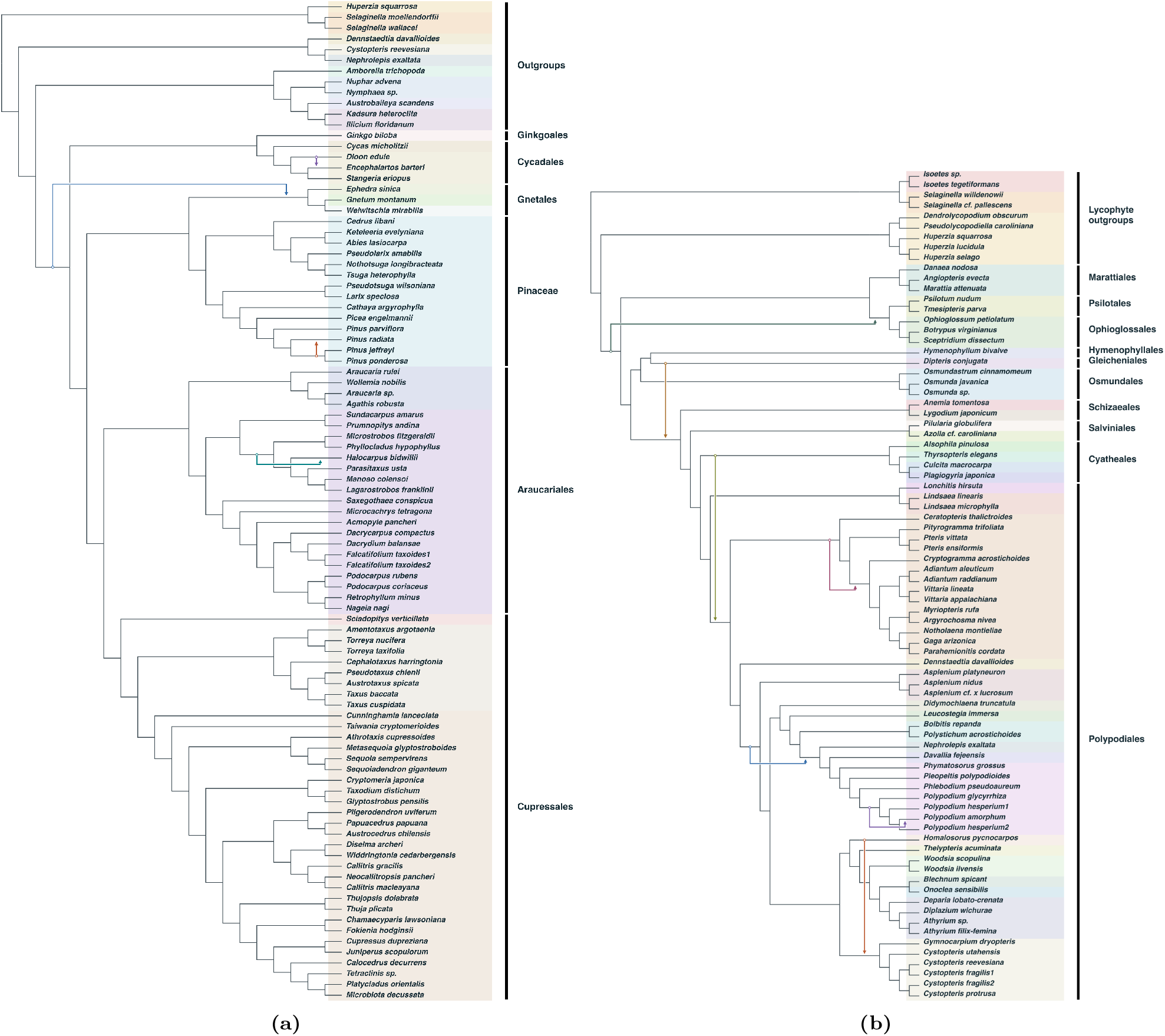
Networks inferred by BROOQS on the gymnosperm and fern subsets of the 1KP dataset. (a) Gymnosperm subset. (b) Fern subset.

**Figure S9:**
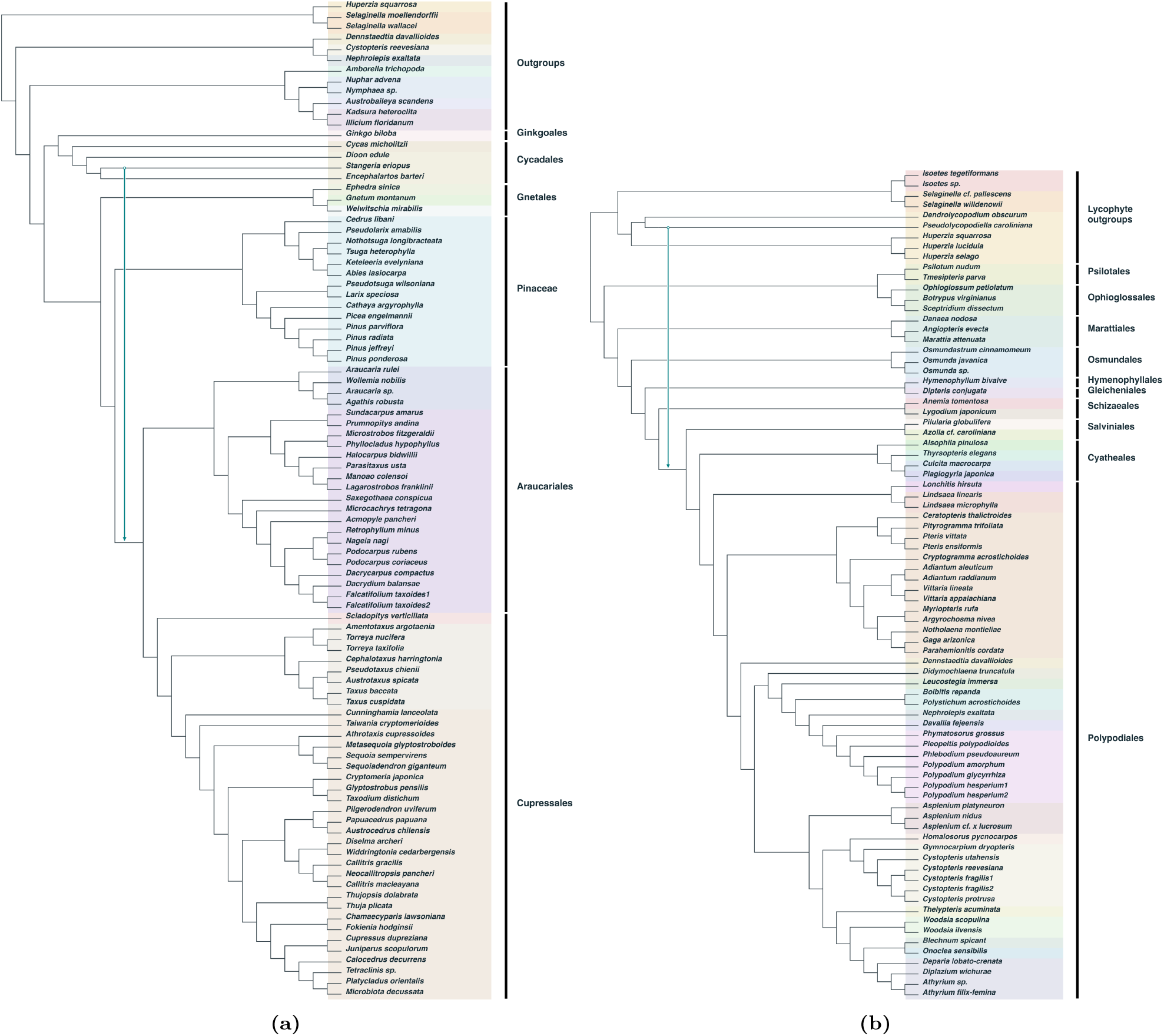
Networks inferred by CAMUS on the gymnosperm and fern subsets of the 1KP dataset. (a) Gymnosperm subset. (b) Fern subset. Reticulations inferred within the outgroups are not presented.

**Figure S10:**
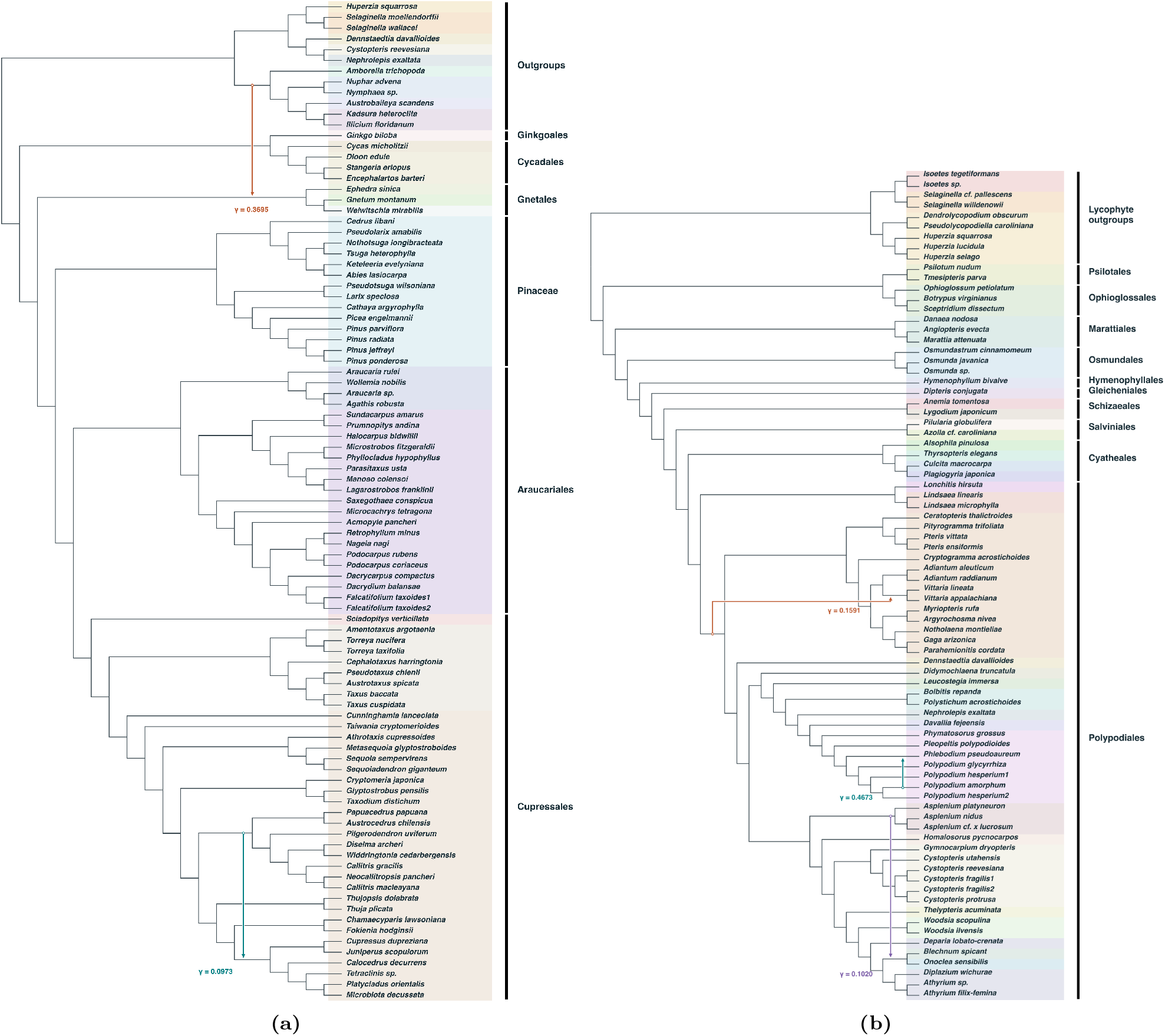
Networks inferred by InPhyNet on the gymnosperm and fern subsets of the 1KP dataset. (a) Gymnosperm subset; reticulations inferred within the outgroups are not presented. (b) Fern subset.

**Figure S11:**
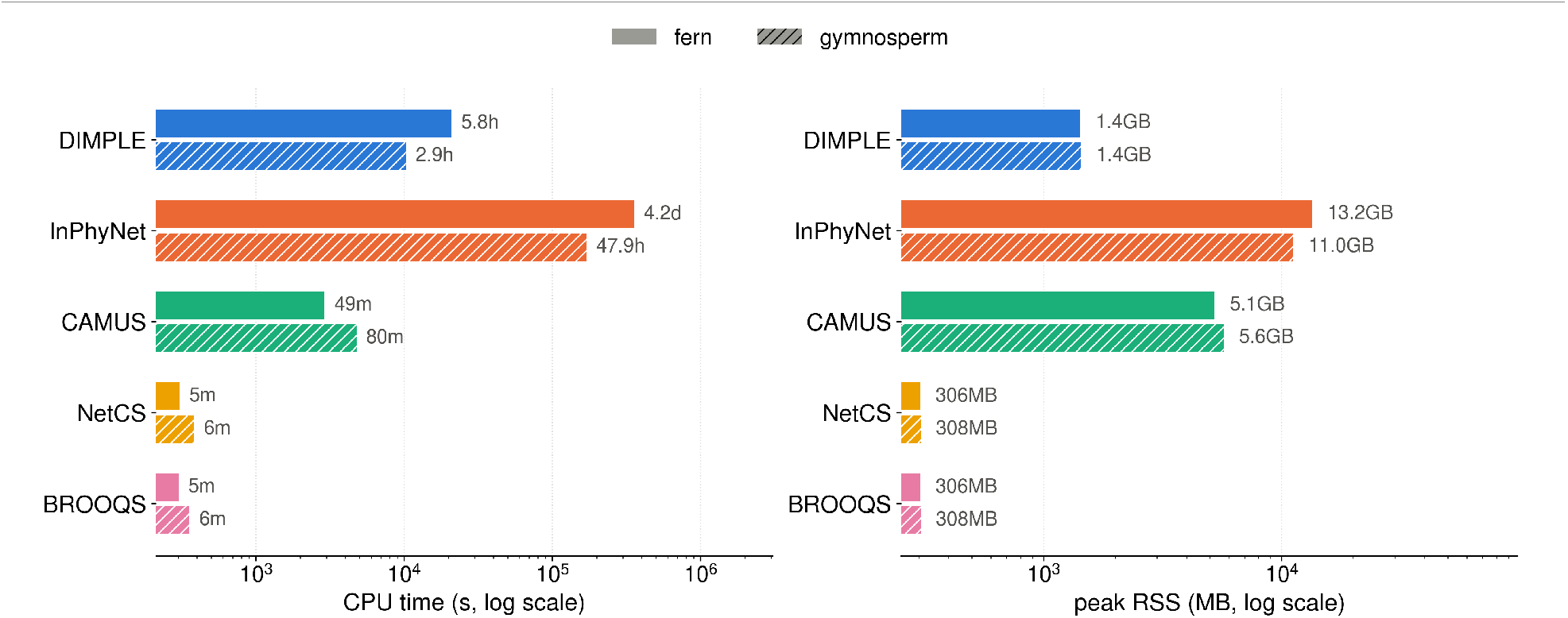
Runtime and memory of DIMPLE versus other methods on the two empirical datasets. Here InPhyNet timing excludes reticulation-count selection because SNaQ was run at a single reticulation count per subset. We used the taxon subsets reported in the InPhyNet paper’s supplementary information; these reach 21 taxa and dominate the computation time. However, for DIMPLE, we include MPL runtime for each of the reticulation counts 0, 1, 2, and 3, which we used for reticulation-count selection. Left: total CPU time. Right: peak resident memory. MPL inference accounts for over 96% of DIMPLE’s CPU time.

